# Tropical tree growth sensitivity to climate is driven by species intrinsic growth rate and leaf traits

**DOI:** 10.1101/2021.06.08.447571

**Authors:** David Bauman, Claire Fortunel, Lucas A. Cernusak, Lisa P. Bentley, Sean M. McMahon, Sami W. Rifai, Jesús Aguirre-Gutiérrez, Imma Oliveras, Matt Bradford, Susan G. W. Laurance, Guillaume Delhaye, Michael F. Hutchinson, Raymond Dempsey, Brandon E. McNellis, Paul E. Santos-Andrade, Hugo R. Ninantay-Rivera, Jimmy R. Chambi Paucar, Oliver L. Phillips, Yadvinder Malhi

**Affiliations:** Environmental Change Institute, School of Geography and the Environment, University of Oxford, Oxford, UK; Smithsonian Environmental Research Center, Edgewater, Maryland 21037, USA; AMAP (Botanique et Modélisation de l’Architecture des Plantes et des Végétations), Université de Montpellier, CIRAD, CNRS, INRAE, IRD, Montpellier, France; Centre for Tropical Environmental and Sustainability Science, College of Science and Engineering, James Cook University, Cairns, Queensland 4878, Australia; Department of Biology, Sonoma State University, 1801 E. Cotati Ave., Rohnert Park, CA 94928 USA; ARC Centre of Excellence for Climate Extremes, University of New South Wales, Sydney, NSW, Australia; Biodiversity Dynamics, Naturalis Biodiversity Center, Leiden, The Netherlands; CSIRO Land and Water, Tropical Forest Research Centre, Atherton, Queensland, Australia; Fenner School of Environment and Society, The Australian National University, Canberra, Australia; Department of Plant and Environmental Sciences. New Mexico State University. Las Cruces, NM, USA; Universidad Nacional San Antonio Abad del Cusco, Cusco, Perú; School of Geography, University of Leeds, Leeds, UK

**Keywords:** tropical forest ecology, demography, tree vital rates, functional traits, climate change, vapour pressure deficit (VPD), climate anomalies, water use efficiency, photosynthesis, permanent plots

## Abstract

A better understanding of how climate affects growth in tree species is essential for improved predictions of forest dynamics under climate change. Long-term climate averages (mean climate) and short-term deviations from these averages (anomalies) both influence tree growth, but the rarity of long-term data integrating climatic gradients with tree censuses has so far limited our understanding of their respective role, especially in tropical systems. Here, we combined 49 years of growth data for 509 tree species across 23 tropical rainforest plots along a climatic gradient to examine how tree growth responds to both climate means and anomalies, and how species functional traits mediate these tree growth responses to climate. We showed that short-term, anomalous increases in atmospheric evaporative demand and solar radiation consistently reduced tree growth. Drier forests and fast-growing species were more sensitive to water stress anomalies. In addition, species traits related to water use and photosynthesis partly explained differences in growth sensitivity to both long-term and short-term climate variations. Our study demonstrates that both climate means and anomalies shape tree growth in tropical forests, and that species traits can be leveraged to understand these demographic responses to climate change, offering a promising way forward to forecast tropical forest dynamics under different climate trajectories.

## Introduction

Tropical forests are key contributors to global carbon sequestration (Pan *et al*. 2011; Needham *et al*. 2018), but climate change may reduce this important ecosystem service by suppressing tree growth or increasing mortality risk, particularly in warmer and drier tropical forests (Brodribb *et al*. 2020; Sullivan *et al*. 2020). Hence it is important to understand how climate influences tree growth, both through long-term local averages (hereafter, “mean climate”, often calculated over a period of 30 years) and short-term deviations from these averages (hereafter, anomalies, estimated as the difference from a 30-year baseline average for a particular place and time (Rifai *et al*. 2018, 2019). Long-term mean climate can constrain the ways species achieve different growth rates in different locations through their effect on tree physiological processes (Rifai *et al*. 2018; Green *et al*. 2019; Sullivan *et al*. 2020). On the other hand, climate change manifests in particular as increases in the magnitude of anomalies and frequency of ‘extreme weather events’ (i.e. extreme anomalies) (Jentsch *et al*. 2007; Malhi *et al*. 2009; Harris *et al*. 2018), which can alter growth rates at the scale of weeks, months, and years (Mendivelso *et al*. 2014; Rifai *et al*. 2018, 2019; Sanginés de Cárcer *et al*. 2018; Yuan *et al*. 2019; Grossiord *et al*. 2020). So far, most studies on the impact of anthropogenic climate change on species and community growth rates have focused either on mean climate or extreme climatic events (e.g. Phillips *et al*. 2009; Fadrique *et al*. 2018; Aguirre-Gutiérrez *et al*. 2019). However, predicting tropical forest dynamics requires disentangling the relative effects of long-term mean climate and the continuum of small to large climate anomalies on tree growth (Harris *et al*. 2018).

How species differences influence tree growth response to mean climate and climate anomalies remains unclear. Functional traits (*sensu* Violle et al., 2007) can capture species differences in ecological strategies and allocation tradeoffs to growth, survival and reproduction (Westoby *et al*. 2002; McGill *et al*. 2006). Trait-based approaches offer a path towards a more mechanistic understanding of species differences in tree growth response to climate drivers (Wagner *et al*. 2014; Uriarte *et al*. 2016; Brodribb *et al*. 2020; Laughlin *et al*. 2020). Specifically, the ‘fast-slow’ plant economics spectrum links fast-growing and slow-growing species to acquisitive and conservative trait values, respectively (Reich 2014). As high growth rates may come with a cost of lower stress-tolerance (Reich 2014; Gibert *et al*. 2016), acquisitive strategies could increase growth sensitivity to climate anomalies, while conservative strategies could attenuate it. Physiological traits directly related to photosynthesis and water use efficiency are good candidates to reflect the effects of light- and water-related climate variables on tree growth and forest dynamics (Wagner *et al*. 2014; Brodribb *et al*. 2020; Powers *et al*. 2020; Rowland *et al*. 2021).

Uncoupling mean climate and climate anomalies as drivers of tree growth and understanding the functional basis for differences in species growth responses to climate requires detailed long-term inventories stretching along climatic gradients, coupled with information on species traits. However, most studies have focused on a single site (Condit *et al*. 2017), growth-climate relations (Rifai *et al*. 2018), growth-trait relations (Poorter *et al*. 2008; Paine *et al*. 2015; Gibert *et al*. 2016; Gray *et al*. 2019) or trait-environment relationships along climatic gradients (Aguirre-Gutiérrez *et al*. 2019; Rosas *et al*. 2019), with few studies combining all these aspects (Fyllas *et al*. 2017). Here, we take advantage of a unique 49-year dataset of regularly-censused tropical tree growth (two to five year-intervals) spanning 509 species across 23 plots covering an elevation gradient of 1300 m and encompassing a broad range of climatic conditions, in North Queensland (Wet Tropics of Australia). We use 15 morphological, chemical and physiological traits related to leaf, wood and maximum size collected within the plot network for 75 dominant species to test how these traits mediate species growth responses to climate drivers. We couple the multi-year census data with the detailed plant traits dataset in Bayesian hierarchical models to relate tree growth, species traits, forest plots, and climatic data (Fig. 1). We examine the effects of both mean climate and climate anomalies on interannual tree growth variation, both within and across species, and evaluate the role of functional traits in capturing species differences in growth sensitivity. We also test whether the effects of climate anomalies on plot-level growth rate variation depend upon long-term mean climate. Specifically, we ask:

i. How do mean climate and climate anomalies determine interannual variation in tree growth rates, and what are the main climatic drivers?
ii. Are drier and warmer forests more sensitive to positive anomalies in temperature and water stress?
iii. Can functional traits explain interspecific differences in growth sensitivities to climate?

**Figure 1:**
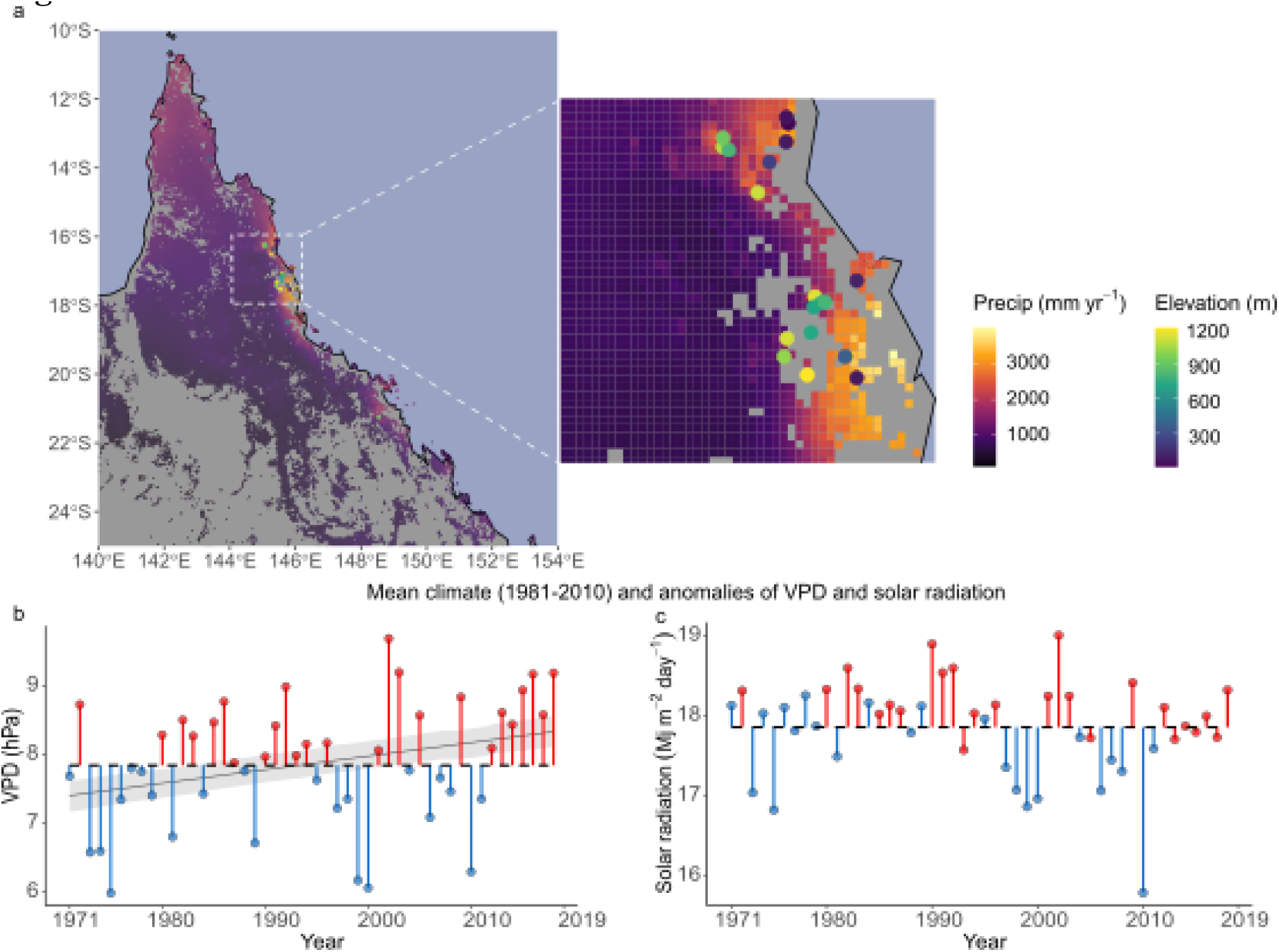
Spatial and temporal dimensions of the tropical forest network. **a**: Maps of North Queensland (Australia) and the 23 forest plots on a background of the long-term mean annual precipitation for woody vegetation areas. Circles: plots; Circle colours: Plot elevation (strongly negatively correlated to mean annual temperature). **b** and **c**: Illustration of the temporal extent of the study and of the concepts of mean climate and anomalies for one plot (Mont Haig) presenting vapour pressure deficit (VPD) and solar radiation (SRAD) through time, respectively. Fig.1b,c show the mean climate (1981-2010) (horizontal black dashed line) and negative and positive anomalies (blue and red vertical segments and dots; monthly anomalies averaged per year). VPD and SRAD were modelled as a plot-specific function of year (see *Methods* and Table S4). The thin black line and shaded areas are the median and 95%-highest posterior density interval (HPDI) of the slope characterising the VPD increase over time. SRAD did not present any clear trend (slope not represented; i.e. the 95%-HPDI encompassed zero).

## Materials and Methods

### Study sites and demographic data

Individual tree annual absolute growth rates were calculated for 12,853 trees in 23 permanent forest plots of tropical rainforest located in northern Queensland, Australia, between 12°44’ S to 21°15’ S and 143°15’ E to 148°33’ E, and encompassing an elevation gradient between 15 and 1200 m a.s.l. and a period of 49 years (Fig. 1a; Table S1) (20 CSIRO long-term plots (Bradford *et al*. 2014), and three more recent plots; see Supplementary Methods S1). Regular cyclonic disturbance contributes to the dynamics of the forests (Murphy *et al*. 2013). They cover a wide range of mean annual temperatures (19°C to 26.1°C), precipitations (1213 to 3563 mm), solar radiation (17.8 to 19.4 Mj m^-2^ day^-1^) and vapour pressure deficit (6.5 to 11.8 hPa) (Table S1). At plot establishment, all trees with stems ≥ 10 cm diameter at breast height (DBH) were mapped, identified to species level and measured for diameter. The 20 long-term plots were re-measured every two years for ten years, and then at three- to four-year intervals, with diameter, recruits and deaths recorded, summing up to 11 to 17 censuses per plot. The remaining three plots were established between 2001 and 2012 and resampled one to three times (Table S1).

All available censuses were used to calculate individual annualised absolute growth rate (AGR) based on DBH at date 1 and 2 (*t*_1_ and *t*_2_), as:

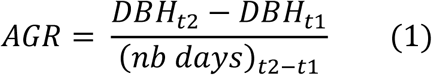

Abnormal AGR values were removed following (Condit *et al*. 2004) (see Supplementary Methods S1). Pteridophytes and palms species were excluded from the analyses due to their lack of secondary growth.

### Climate data

The effect of climate on growth was studied through four climate variables encompassing a wide range of variability across the plots and relevant for tree growth (see details in Supplementary Methods S1): mean temperature (Tmean), solar radiation (SRAD), vapour pressure deficit (VPD), and maximum climatological water deficit (MCWD; a proxy of the annual accumulated water stress over the drier season, estimated from climate data as the cumulative deficit between precipitation and evapotranspiration, hence better capturing the seasonality of precipitation and potential soil water deficit than precipitation itself (Aragão *et al*. 2007; Malhi *et al*. 2009, 2015) (Table S1, Table S3a).

Climate data collection is detailed in the Supplementary Methods S1 and summarised here. Monthly climatic variables were obtained for the period 1970 to 2018 for each plot from ANUClimate v.2.0 (Hutchinson *et al*. 2014). The monthly actual evapotranspiration (aet) was derived from TerraClimate (Abatzoglou *et al*. 2018). The aet was used in combination with rainfall to calculate the monthly climatological water deficit (CWD). The CWD was reset to zero at the wettest month of the year and had an upper bound at 1000 mm. It was used to calculate monthly MCWD through a rolling maximum over the previous 12 months.

In each forest plot, a monthly 30-year historical mean and standard deviation were calculated over the 1981-2010 period for Tmean, SRAD, VPD, and MCWD (Table S1). On this basis, we calculated in each plot the monthly anomalies for each variable (i.e. monthly 30-year mean *μ* subtracted from monthly value) and divided them by their location-specific 30-year monthly standard deviation *σ*, yielding standardised anomalies (Aragão *et al*. 2007; Rifai *et al*. 2018):

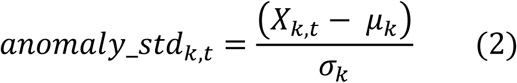

where *X_k, t_* is the climate variable value in plot *k* at time *t* (i.e. year and month), and *μ_k_* and *σ_k_* are the monthly 30-year mean and standard deviation of the corresponding plot’s location.

Standardised anomalies are expressed in units of standard deviation from monthly means over 1981-2010. This allows the comparison of plots differing not only in their historical means but also in the long-term variation range around them, that is, an important element to detect anomaly effects on tree growth across different mean climates.

To build the climate covariates for the tree growth models, the monthly 30-year mean and standardised anomaly variables were averaged over the months between consecutive censuses (two to five years). For MCWD, the maximum over the growth periods between two censuses was used instead of the weighted mean. The eight resulting interannual averaged variables were used as predictors to model tree growth (see *Data analysis*). Correlations among these variables, stand structure and elevation are presented in Table S3a and the Supplementary Methods S1.

### Stand structure

As stand structure can vary between plots, we include its effect on tree growth through total plot basal area. Plot basal area (m²/ha) was calculated at each census, with expectations that increasing basal area would have a general negative effect on tree growth (Sánchez-Salguero *et al*. 2015; Muledi *et al*. 2020).

### Functional traits

Between July and September 2015, we measured 15 traits of 75 dominant, canopy tree species in eight plots along the gradient (Table 1; Table S1 and S2 for plot and species details). Species were chosen to sample those that made up 80% of the standing biomass. Trait data collection and measurement are detailed in Supplementary Methods S1. We measured leaf, wood and maximum size traits that relate to light, water and nutrient use (Table 1, see Table S3b for pairwise trait correlations, and Fig. S1 for trait distribution along the elevation gradient). Traits were measured on three individuals per species, and included photosynthesis and stomatal conductance at a reference CO_2_ concentration of 400 µmol mol^-1^ and irradiance of 1500 µmol photons m^-2^ s^-1^ (Asat and gsat), dark respiration (Rd) at the same CO_2_ concentration, the CO_2_-saturated photosynthesis and stomatal conductance (Amax and gmax), measured at 1200 µmol mol^-1^ CO_2_. The one-point method (De Kauwe *et al*. 2016) was used to estimate maximum carboxylation rate (Vcmax) for each individual from net photosynthesis measured at 400 µmol mol^-1^ CO_2_, and maximum light-driven electron flux (Jmax) from net photosynthesis measured at 1200 µmol mol-1 CO_2_ (Bloomfield *et al*. 2018). We also measured leaf stable carbon isotope ratio (δ^13^C), nutrient concentration, and leaf area, leaf mass per area (LMA), leaf thickness, wood density (from branches, after bark removal). All traits were averaged at the species level for tree growth analyses.

**Table 1:** Functional traits measured and their functions. . These traits were measured on three adult individuals of 75 tree species and used to model intrinsic growth rate and growth response to mean climate, climatic anomalies, and stand structure.

| Organ | Trait type | Trait | Abbreviation | Units | Mean (min, max) | CV (%) | Functional role |
| --- | --- | --- | --- | --- | --- | --- | --- |
| Leaf | Physiology | Net photosynthetic rate at saturating irradiance and ambient CO<sub>2</sub> (400 ppm) | Asat | μmol CO<sub>2</sub> m<sup>-2</sup> s<sup>-1</sup> | 5.44 (0.98, 9.36) | 28.6 | Photosynthesis and growth |
|  |  | Net photosynthetic rate at saturating irradiance and saturated CO<sub>2</sub> (1200 ppm) | Amax | μmol CO<sub>2</sub> m<sup>-2</sup> s<sup>-1</sup> | 12.9 (7.7, 19.2) | 19.1 | Photosynthesis and growth |
|  |  | Stomatal conductance at saturating irradiance and ambient CO<sub>2</sub> (400 ppm) | g<sub>sat</sub> | mol H<sub>2</sub>O m<sup>-2</sup> s<sup>-1</sup> | 0.071 (0.02, 0.146) | 34.8 | Control of carbon and water exchange between the leaf and the atmosphere, hence influencing photosynthesis and water use efficiency (Liu *et al.* 2017) |
|  |  | Stomatal conductance at saturating irradiance and saturated CO<sub>2</sub> (1200 ppm) | g<sub>max</sub> | mol H<sub>2</sub>O m<sup>-2</sup> s<sup>-1</sup> | 0.064 (0.018, 0.135) | 33.1 | Control of carbon and water exchange between the leaf and the atmosphere, hence influencing photosynthesis and water use efficiency (Liu *et al.* 2017) |
|  |  | Maximum rate of electron transport | Jmax | μmol e<sup>-</sup> m<sup>-2</sup> s<sup>-1</sup> | 68.2 (39.3, 94.5) | 18.0 | Directly related to photosynthetic rate (Walker *et al.* 2014) |
|  |  | Maximum rate of Rubisco carboxylation | Vcmax | μmol CO<sub>2</sub> m<sup>-2</sup> s<sup>-1</sup> | 31.2 (11.2, 52.1) | 25.3 | Directly related to photosynthetic rate s |
|  |  | Ratio of maximum electron transport to maximum carboxylation rates | Jmax / Vcmax | μmol e<sup>-</sup> μmol<sup>-1</sup> CO<sub>2</sub> | 2.32 (1.67, 6.73) | 26.9 | Relative allocation to Jmax and Vcmax (Smith *et al.* 2019) |
| Leaf | Metabolism | Maximum rate of dark respiration | Rd | μmol CO<sub>2</sub> m<sup>-2</sup> s<sup>-1</sup> | 0.826 (0.259, 1.74) | 41.7 | Metabolic rate; Correlates with photosynthetic capacity (Atkin *et al.* 2015) |
| Leaf | Chemistry | Leaf carbon stable isotope ratio | leaf δ<sup>13</sup>C | ‰ | -30.4 (-32.9, -27.7) | 4.6 | Positively correlated with intrinsic water use efficiency and the ratio of intercellular to ambient CO<sub>2</sub> concentrations, hence relying on stomatal conductance and photosynthetic capacity (Cernusak *et al.* 2013) |
|  |  | Leaf nitrogen per unit area | N<sub>leaf</sub> | μg cm<sup>-2</sup> | 177 (104, 268) | 22.6 | N<sub>leaf</sub> mainly supports the photosynthetic machinery, mostly the Rubisco carboxylation rate and hence photosynthesis (Wright *et al.* 2004; Walker *et al.* 2014; Quebbeman & Ramirez 2016) |
|  |  | Leaf phosphorus per unit area | P<sub>leaf</sub> | μg cm<sup>-2</sup> | 10.3 (4, 38.7) | 46.5 | Important determinant of photosynthetic rate (Walker *et al.* 2014) |
| Leaf | Structure | Leaf area | LA | cm<sup>2</sup> | 540 (7, 32864) | 695.9 | Light capture efficiency and control of the boundary layer driving leaf heating-cooling dynamics (Wright *et al.* 2017) |
|  |  | Leaf thickness | thickness | mm | 0.28 (0.17, 0.53) | 25.6 | Increases structural support and leaf lifespan; Related to resource acquisition and use (Westoby *et al.* 2002; Vile *et al.* 2005) |
|  |  | Leaf mass per area | LMA | g m<sup>-2</sup> | 126 (78, 216) | 27.4 | Relative allocation to biomass per leaf area; Strongly correlated to leaf lifespan and thus nutrient use efficiency (Osnas *et al.* 2013) |
| Wood | Structure | Wood density | WD | g cm<sup>-3</sup> | 0.585 (0.312, 0.795) | 17.1 | Mechanical support, water transport and storage capacity (carbon and other nutrients, defence compounds) (Chave *et al.* 2009) |
| Whole plant | Maximum size | Maximum diameter (130 cm) | DBH<sub>max</sub> | cm | 54.6 (16.4, 113.2) | 41.6 | Proxy of maximum height, itself summarising light-acquisition and growth strategies (Westoby 1998; Rüger *et al.* 2012) |

### Data analysis

We addressed our questions through three sets of Bayesian multilevel models (M1 to M3; details in Supplementary Methods S1).

#### M1: Tree growth response to climate means and anomalies, and species differences in their sensitivities to climate

In M1, we used 12,853 individuals from all 509 species to test the effects of climate on tree growth, and to investigate tradeoffs among species between intrinsic growth rate and growth sensitivity to climate covariates. We built a two-level hierarchical Bayesian model of AGR, where the hierarchy included an upper level of response (hereafter grand coefficients or effects, affecting AGR across species) above a lower, species-level response. The higher level modelled AGR responses to covariates via hyperparameters (i.e. statistical distributions from which species-level intercepts and slope coefficients arose), while the lower level captured species-specific growth sensitivities to model covariates, and species-level intercepts (hereafter intrinsic AGR) captured unexplained growth variation across individuals, growth periods, and plots.

More specifically, we modelled individual log(AGR) as a species-specific function of (i) initial tree size (approximated by log(DBH) at the beginning of a growth period), (ii) the local 30-year mean of a climate variable, (iii) the anomalies of the same climate variable averaged over the studied growth period, and (iv) stand structure (approximated by plot basal area at the beginning of a growth period), using varying slopes (also known as random slopes) and a covariance matrix to estimate correlations among species-specific AGR sensitivities to the covariates, as:

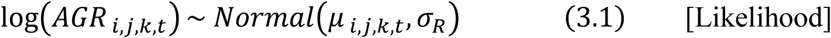

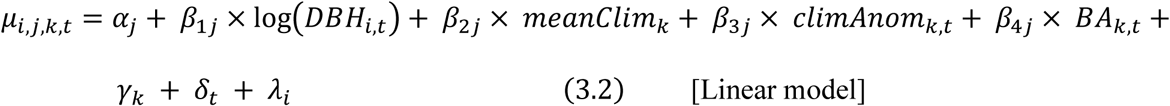

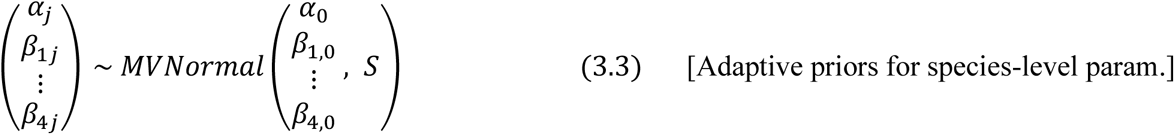

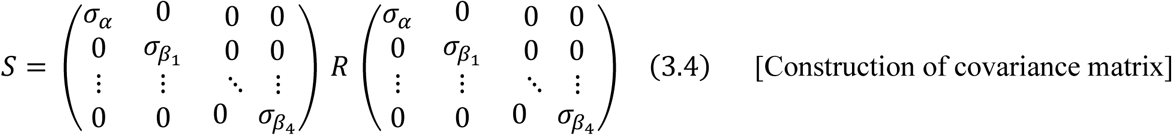

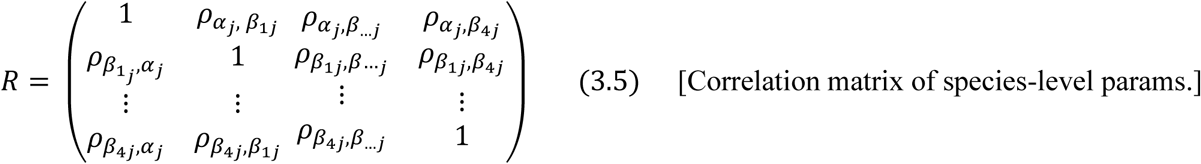

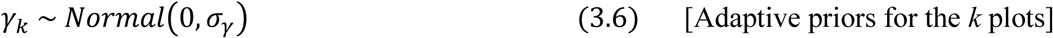

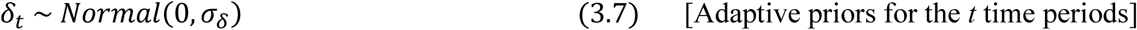

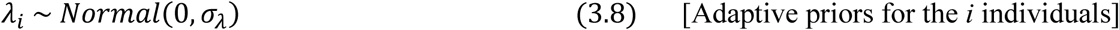

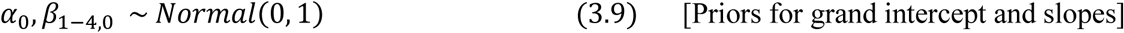

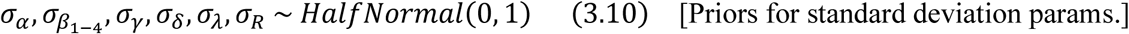

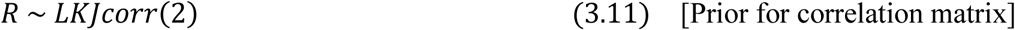

where *α_j_* characterises the intrinsic AGR of species *j* and *β*_1*j*_, *β*_2*j*_, *β*_3*j*_ and *β*_4*j*_ characterise the AGR response of species *j* to tree size, mean climate (1981-2010), standardised climate anomalies and plot basal area in plot *k* for time interval *t*. The parameter *α_0_* represents the grand intercept, and the parameters *β_1-4, 0_* are the grand slopes of model covariates whose posterior distributions represent the effect of mean climate and climate anomaly on AGR across species.

The matrix of fitted correlation coefficients among species-level intercepts and slopes (*α_j_, β*_1*j*_, *β*_2*j*_, *β*_3*j*_ and *β*_4*j*_) allows evaluating correlations among species intrinsic growth rate (intercepts *α_j_*) and species AGR sensitivity to model covariates (*β_1-4j_*). For instance, a model with a negative *ρ_αj_*_, *β*3*j*_ parameter and a negative *β*_3,0_ slope would indicate that species with higher intrinsic growth rate (*α_j_*) tend to have higher sensitivity (i.e. more negative slopes) to climate anomalies. Using covariance matrix to pull information across species-level intercepts and slopes through the multinormal distribution improves the accuracy of posterior likelihood estimates both across and within species (hierarchical levels 1 and 2, respectively) while limiting risks of overfitting through adaptive regularising priors, or partial pooling (e.g. McElreath 2020).

Parameters *γ_k_*, *δ_t_*, *λ_i_* are varying intercepts capturing the residual variation in expected individual AGR occurring among forest plots, time periods between consecutive censuses (characterised by the years beginning and ending a given census period), and individual stems, respectively. This model was run separately for each of the four climate variables (Tmean, SRAD, VPD, and MCWD) to manage model complexity (representing a total of four M1 models).

#### M2: Trait-mediated species-level tree growth response to climate

Models M2 have the same hierarchical structure as M1, but additionally include the role of species traits in AGR response to climate. We thus used a subset of 5,191 individuals from the 75 species with trait data. In M2, the species-level intercept and slopes are modelled as depending from species mean trait value such that both species-specific intrinsic AGR and AGR sensitivity to covariates can be influenced (either accentuated or lessened) by species traits (Rüger *et al*. 2012; Uriarte *et al*. 2016; Fortunel *et al*. 2018) as:

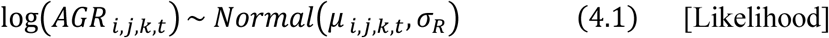

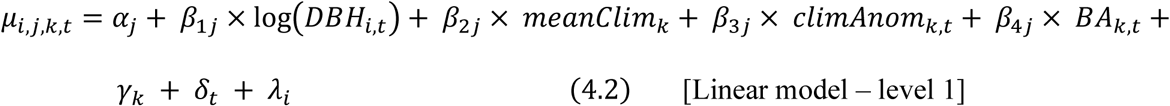

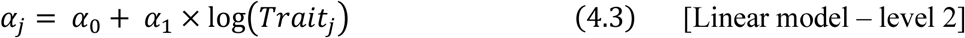

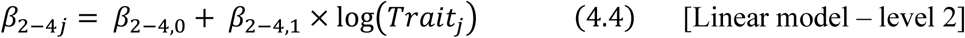

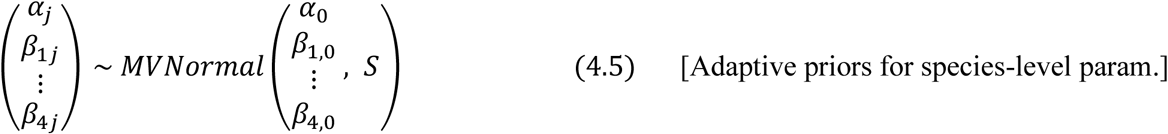

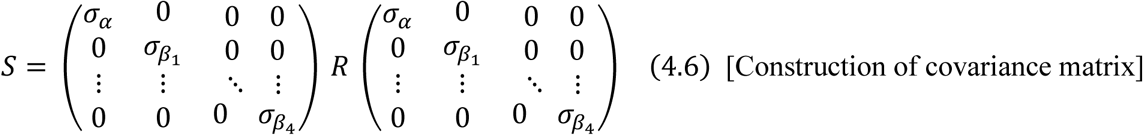

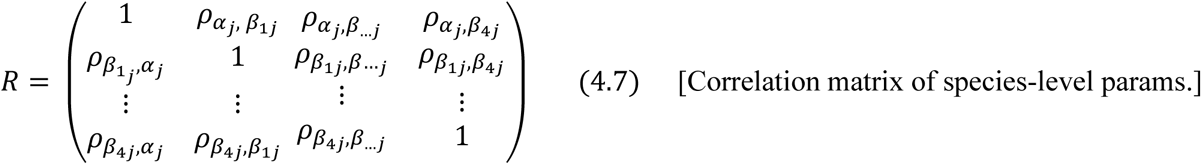

where eqs. 4.1, 4.2, 4.5-4.7 are the same as eqs. 3.1-3.5 of M1, while species-level intercepts and slopes are mediated by species mean trait value (eqs. 4.3-4.4; detailed equations and priors in Supplementary Methods S1). Parameter *α_1_* is the species-level departure from the grand intercept (*α_0_*) for an increase of one standard deviation in the log(trait T*_j_*) value of species *j* (direct effect of trait on AGR), while *β*_2-4, *1*_ are the departures from the grand slope of the corresponding model covariates for an increase of one standard deviation in the log(trait T*_j_*) value of species *j* (trait mediation of AGR response to climate and stand structure; see Supplementary Methods S1 for ecological interpretations of trait coefficient signs). We did not include the role of species traits in AGR response to tree size because some traits can change through tree ontogeny (Fortunel *et al*. 2020) and our trait data does not encompass species tree size ranges. M2 models were run separately for each of the four climate variables and for each of the 15 functional traits to manage model complexity (representing a total of 60 M2 models).

In both M1 and M2 models, we standardised the response variable log(AGR) and all covariates – but climate anomalies – to mean zero and unit standard deviation, to allow relative importance comparisons between covariates through slope coefficients (Schielzeth 2010), and to ease plausible weakly-informative prior assignment to the parameters (McElreath 2020) (see Supplementary Methods S1). We did not standardise averaged monthly anomalies to maintain their interpretability as deviations from long-term means in terms of plot-specific units of standard deviation (see eq. 2; i.e. mean anomaly covariate slope coefficients are not directly comparable to other covariate mean slopes).

#### M3: Plot-level tree growth response to climate anomalies and interaction with mean climate

M3 models evaluate plot-level growth response to climate anomalies, and whether it varies depending on local mean climates (e.g. whether plot-level AGR sensitivity to VPD anomalies is higher in drier sites). We focused on the tree growth at the plot level, and modelled the expected log(AGR) as a linear function of mean climate and climate anomalies. We used a similar Bayesian hierarchical model as described for M2, where plot-specific average AGR depended on climate anomalies, whose effect on AGR itself depended on the plot mean climate, as:

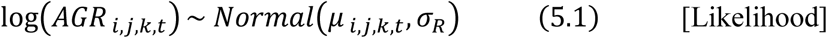

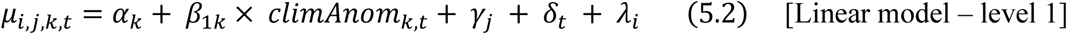

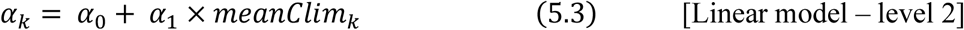

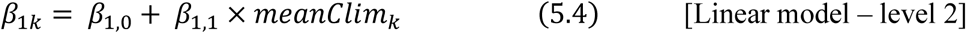

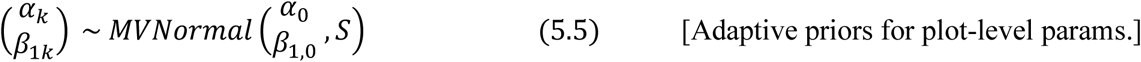

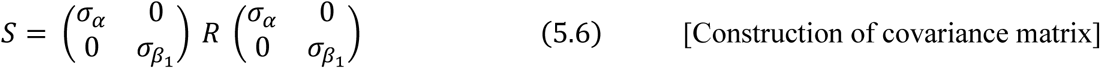

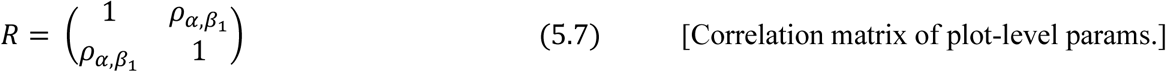

where *α_k_* is the average growth rate in plot *k*, and *β_1k_* characterises the growth response of plot *k* to standardised climate anomalies for time interval *t*. *α_0_* is the mean intercept value (i.e. mean absolute growth rate) across plots, and *α_1_* is the departure from the grand mean for one unit increase in mean climate (see detailed equations and priors in Supplementary Methods S1). *β_1,0_* is the grand slope of climate anomalies, and *β_1,1_* is the departure from this grand mean for a one unit increase in mean climate (mediation of the effect of anomalies on growth by the plot mean climate). Parameters *γ_j_*, *δ_t_*, *λ_i_* are varying intercepts for species, census periods, and individual stems, respectively.

We run M3 models only for two climate variables (VPD and SRAD), as we found they were the most important climate variables for tree growth in M1 and M2 models (see *Results*). Standardisation of variables was carried out as for M1.

#### Trends in climate over time

To explore the implications of the effects of climate anomalies on tree growth, we built a separate set of hierarchical Bayesian models to test for linear temporal trends in mean annual climate variables between 1971 and 2019. We used varying *year* slopes per plots to allow plot-specific trends (model details in Supplementary Methods S1). We also run the models for the period 2000 to 2019 for comparison with recent analyses suggesting an increasing rate of VPD increase over time since the late nineties (Yuan *et al*. 2019). Annual mean temperature and VPD increased of 0.015 °C and 0.02 hPa per year between 1971 and 2019 (R² = 0.97 and 0.84, respectively, Table S4; illustration in Fig. 1b) and of 0.038 °C and 0.045 hPa per year between 2000 and 2019 (R² = 0.98 and 0.81, respectively, Table S4). There was no general temporal trend for MCWD or SRAD (Fig. 1c).

#### Analysis of model outcomes

All model parameter posteriors were summarised through their median and 95%-highest posterior density interval (HPDI) (i.e. the narrowest posterior interval encompassing 95% of the probability mass, corresponding to the coefficient values most consistent with the data; (McElreath 2020)). Model covariates were considered important at two high levels of confidence, when their coefficient had a posterior probability of over 95% or 90% of being either positive or negative (HPDI not encompassing zero).

The goodness-of-fit of the models was assessed through the squared Pearson correlation between the observed AGR and the AGR predicted by the fitted model (R^2^). M1 and M2 models had high explanatory power, with R² of 0.46 and 0.52 on average, respectively. M3 models, with VPD and SRAD as climate variables, had an R² of 0.67 and 0.63, respectively.

Bayesian updating of parameters was performed via the No-U-Turn Sampler (NUTS) in Stan (Carpenter *et al*. 2017), using three chains and 3000 steps (1500 warmings). All models mixed well and converged (Rhat within < 0.01 of 1). Models were run in the R environment (Team 2020) using the packages ‘*brms’* (Bürkner 2017), ‘*tidybayes*’ (Kay 2020) and ‘*tidyverse*’ (Wickham *et al*. 2019).

## Results

### Contribution of climate means and anomalies to tree growth

The main climate drivers affecting tree growth across species were the climate means and anomalies in Tmean, SRAD and VPD (Fig. 2, Fig. S3, Table S5). Tree growth was higher in forests with higher mean Tmean, SRAD and VPD (*β_2j_*: 0.17 [0.08, 0.26], 0.05 [0.02, 0.08], and 0.09 [0.02, 0.17], respectively; median and 95%-HPDI; unless otherwise stated, all intervals are 95%-HPDI). However, tree growth was reduced when forests experienced positive anomalies in temperature, SRAD, and VPD (*β_3j_*: -0.12 [-0.17, -0.07], -0.34 [-0.42, -0.26], and -0.13 [-0.19, - 0.06], respectively). Contrary to our expectation, anomalies in MCWD had no clear effect on tree growth across species (Fig. 2; Fig. S2; Table S5). Tree growth sensitivity to climate, stand structure and tree size varied widely among species (illustration in Fig. S3). Similar results were obtained from the M2 models (subset of 75 species with trait data) (Fig. S5a-d, Table S5), though we no longer detected the effects of temperature anomalies and VPD and solar radiation means in this reduced dataset.

**Figure 2:**
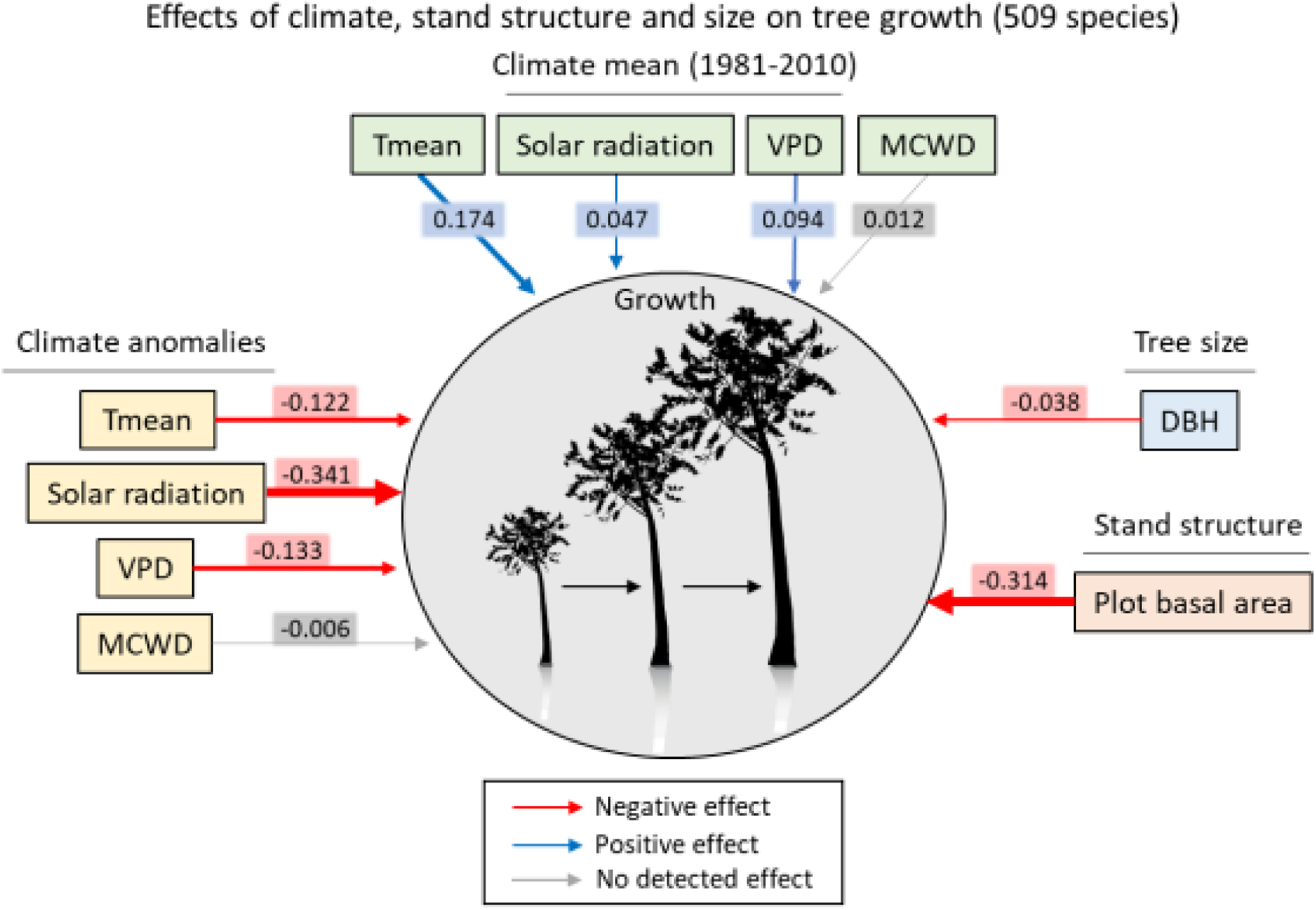
Grand effects of climate, stand structure and tree size on tree growth. (based on all 509 species; M1 models). Red and blue arrows indicate clear negative and positive effects (i.e. slope coefficient 95%-HPDI not encompassing zero). Arrow widths are proportional to the median of covariate slope posteriors (grand slopes, values in rectangles; see *β_1-4,0_* in eqs. 3) (details in Fig. S2 and Table S5).

### Coordinated tree growth sensitivities to climate means and anomalies

Using the fitted matrix of correlations among species-level intercepts and slopes from the M1 models (matrix *R*, see eq. 3.5) allowed testing whether fast- and slow-growing species, and species growing better at opposite extremes of the range of mean climates showed different sensitivities to climate anomalies. Fast-growing species (i.e. with high intrinsic AGR) were more sensitive than slow-growing species to the negative effects of both VPD anomalies and plot basal area on tree growth (Fig. 3c and Fig. S4; *ρ =* -0.36 [-0.48, -0.23] and *ρ =* -0.29 [-0.41, -0.17], respectively). Species that grew better in cloudier forests (i.e. lower SRAD) tended to show steeper growth decreases when experiencing positive anomalies in solar radiation (Fig. 3b; *ρ =* 0.17, [0.01, 0.33]). Species that grew faster in drier forests (i.e. higher VPD) were more negatively affected by positive VPD anomalies (Fig. 3a; *ρ =* -0.15 [-0.29, 0.00]). Finally, species most negatively affected by positive anomalies in VPD also experienced stronger growth decrease in denser forests (high basal area) (Fig. 3d; *ρ =* 0.27 [0.14, 0.40]).

**Figure 3:**
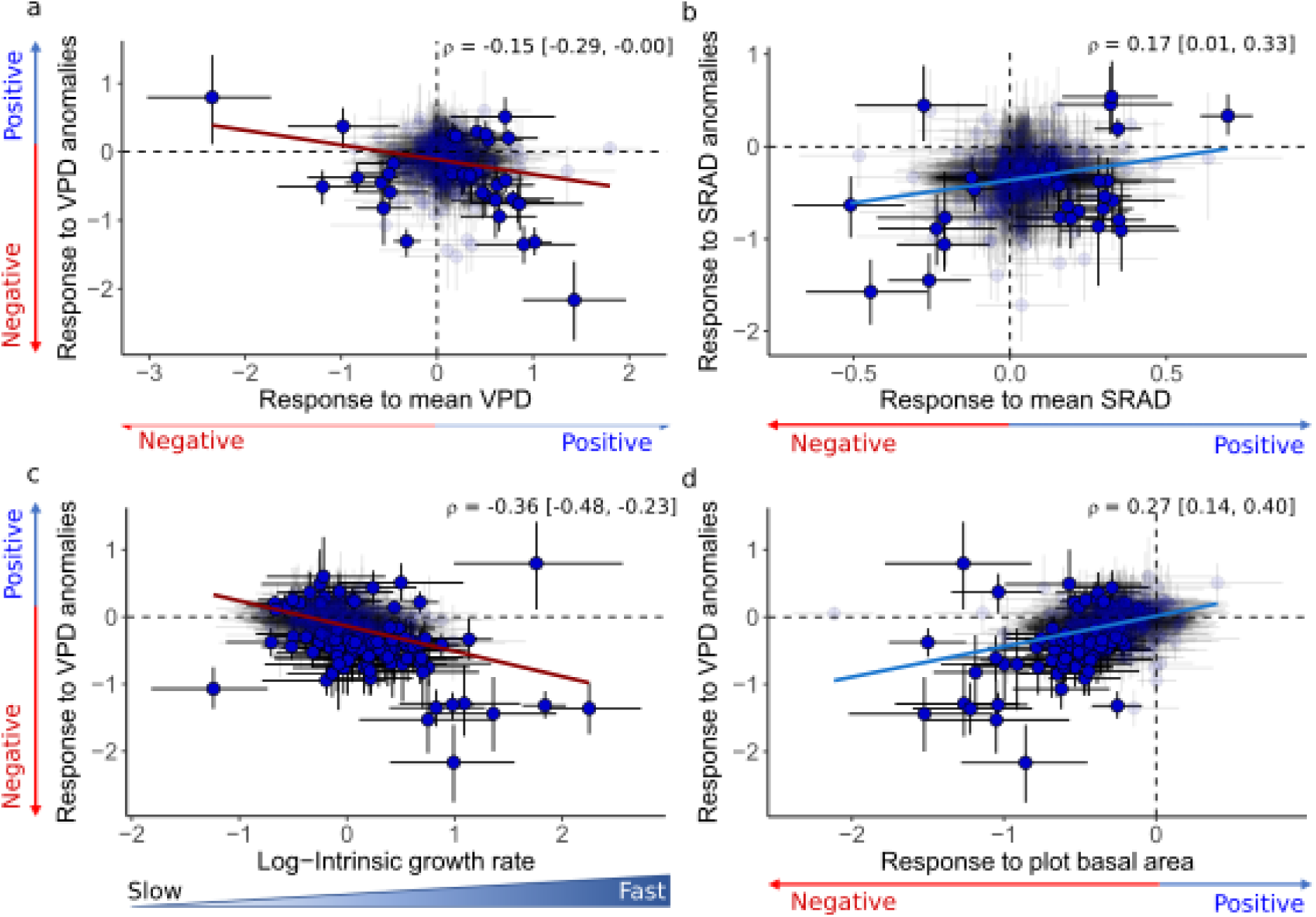
Correlations among species-level growth sensitivities highlighting joint responses to multiple drivers. (M1 models; 509 species). Joint growth sensitivities to: **a**: VPD anomalies and mean VPD; **b**: Solar radiation anomalies and mean solar radiation; **c**: VPD anomalies and intrinsic growth rate; **d**: VPD anomalies and plot basal area. Circles are species, placed at the median of their corresponding coefficient posteriors. Vertical and horizontal bars are 95%-HPDI for the corresponding coefficients. Species for which both plotted coefficients were significant are plain blue; other species are shaded. Blue and red regression lines indicate positive and negative correlations (*ρ*, see eq. 3.5 in Supplementary Methods S1), respectively. Values beyond and below zero indicate positive and negative effects on growth rates, respectively. Mean, lower and upper 95%-HPDI are in the upper right-hand corner of the figures.

### Drier rainforests are more sensitive to VPD anomalies

M3 models highlighted clear interactions between the effects of climate anomalies and mean climate for VPD (*β*_1,1_: -0.26 [-0.39, -0.13]; see eqs. 3), and to a lesser extent for solar radiation (*β*_1,1_: -0.09 [-0.18, -0.01], 90%-HPDI; Table S5). Drier tropical rainforests showed steeper decrease in plot-level growth to positive VPD anomalies (Fig. 4a; Table S5). Cloudier forests exhibited stronger decrease in plot-level growth with positive SRAD anomalies (Fig. 4b; Table S5).

**Figure 4:**
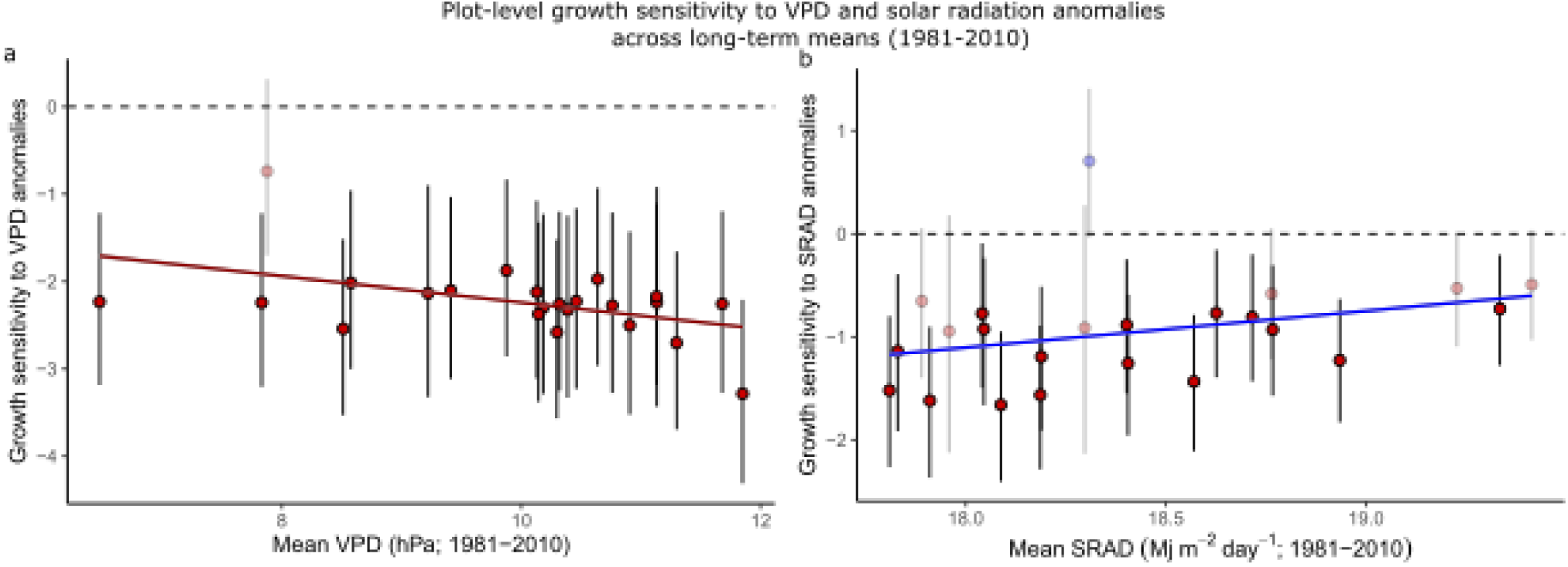
Plot-level growth sensitivity to positive (a) VPD anomalies and (b) solar radiation anomalies (b) across the full range of the corresponding mean climate variable (M3 models). Circles and vertical bars are the median and 95%-HPDI of the plot-level slope posteriors characterising the growth rate responses to climate anomalies. The plot-level models including VPD (a) and SRAD (b) had a marked interaction between anomalies and long-term mean (Table S5), so that plot-level sensitivities to a given anomaly depend on plots’ long-term mean. Figs. 4a and 4b illustrate those interactions through the differences of plot-level growth sensitivity to positive anomalies across the range of long-term means of the corresponding variable. The represented plot-level coefficients were calculated for a positive standardised anomaly equal to the 95th percentile of anomalies in the data, i.e. a standardised anomaly of 0.8 (a) and 0.4 (b). The red and blue regression lines and shaded areas are decreasing and increasing slopes, respectively (median and 95%-HPDI, not encompassing zero), of the represented plot-level coefficients along the long-term means. Horizontal dashed line: limit between positive and negative slope coefficients indicating a growth rate increase and decrease, respectively, with the positive anomaly.

### Functional traits influence species intrinsic tree growth and their response to climate drivers

Based on M2 models, species intrinsic growth increased with dark respiration rate (Rd), DBH_max_, leaf P content, Asat, Vcmax, leaf δ^13^C and LMA. (Fig. 5; Fig. S5e; details in Table S5). Species traits also mediated the effects of climate and forest structure on tree growth, either by accentuating them (species with high values of the trait respond more strongly) or by attenuating them (species with low values of the trait are more sensitive) (Fig. 5; details in Fig. S6 and Table S5). Leaf δ^13^C and P content exacerbated the negative effects of positive anomalies in SRAD on tree growth, while A_max_, g_max_, g_sat_ and J_max_ attenuated them (Fig. 5; Fig. S6f, Table S5). The negative effects of anomalies in VPD on tree growth were exacerbated in species with high leaf δ^13^C, DBH_max_, leaf P, and LMA, further confirming that VPD anomalies had the most negative effects on fast-growing species (Fig. 3c), but also those with low g_max_ or leaf area (Fig. 5; Fig. S6g). Tree growth was less reduced by denser forest environments (high plot basal areas) in species with high wood density, low Rd and low leaf δ^13^C (Fig. 5; Fig. S6i-l).

**Figure 5:**
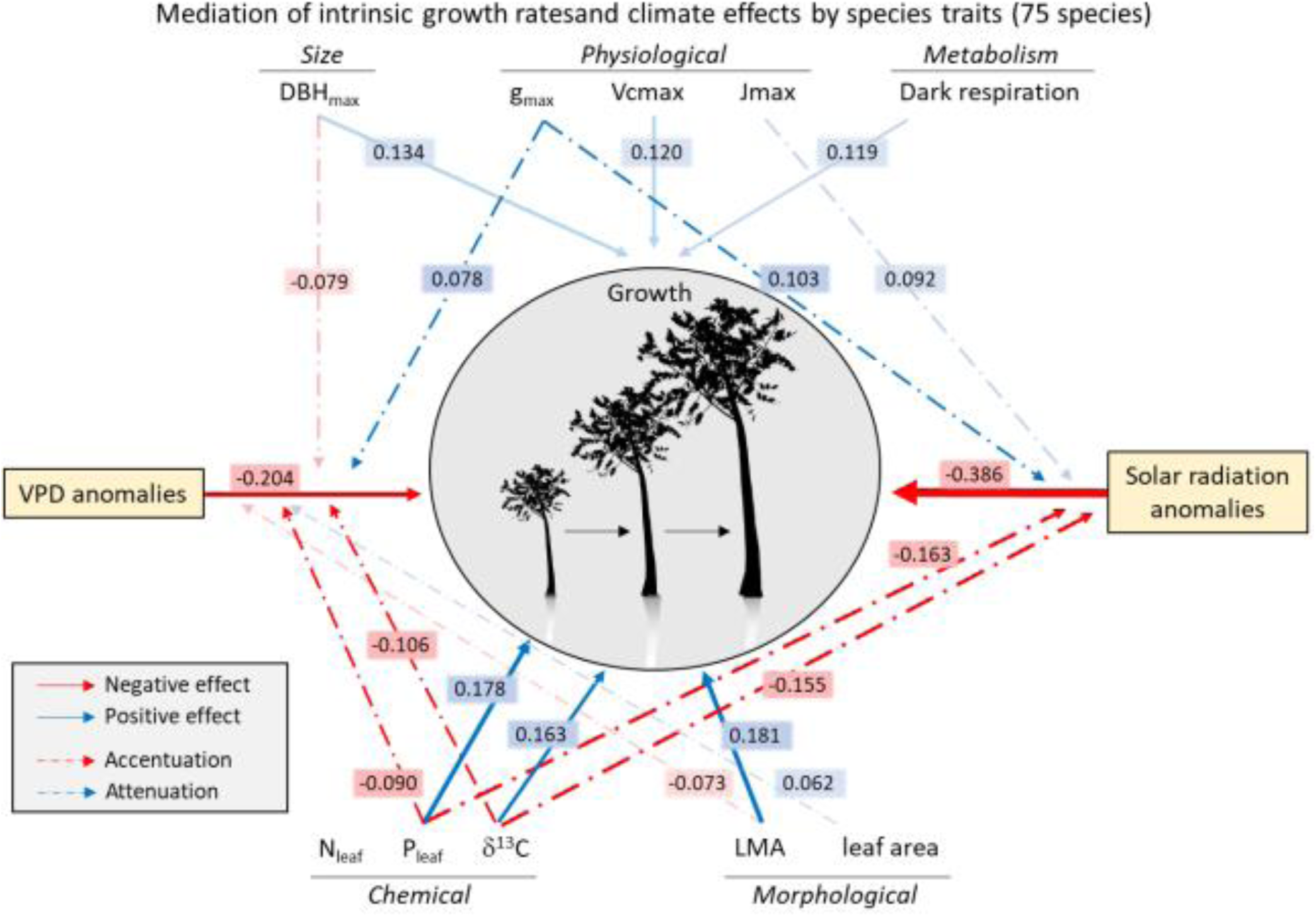
Mediation of intrinsic growth rate and climate anomaly effects on growth rate by species functional traits (M2 models; 75 species). The figure only presents important trait-related effects (95- or 90%-HPDI not encompassing zero; non-transparent and semi-transparent arrows, respectively). Red and blue plain arrows indicate negative and positive direct effects of traits on species’ intrinsic growth rate (*α_1_*, see eq. 4.3). Dashed arrows indicate indirect trait effects on growth through the effects of environmental covariates, i.e. accentuation (red) or attenuation (blue) of the negative effects of VPD or SRAD anomalies when trait values increase (*β_3,1_*, see eq. 4.4). Arrow widths are proportional to the median of the covariate slope posterior across species (i.e. grand slope; details in Fig. S6 and Table S5).

## Discussion

In this study, we disentangled the influences of mean climate and climate anomalies on interannual tree growth and defined how species functional traits mediated climate effects by combining 49 years of demographic data, functional traits and climatic data along a climatic gradient in 23 tropical rainforests of Australia.

### What are the important climatic drivers for tree growth?

Solar radiation (SRAD) and atmospheric water demand (VPD) anomalies were the two overarching climatic drivers of tree growth across pre-existing climatic conditions and species in our study. These two variables were also the main drivers of seasonal stand-level net primary productivity in aseasonal forests across the tropics (Rifai *et al*. 2018), and increasing VPD due to anthropogenic climate change has repeatedly been shown to impact tree growth, biomass and vegetation health (Sanginés de Cárcer *et al*. 2018; Rifai *et al*. 2019; Yuan *et al*. 2019). The pervasive negative effect of VPD anomalies on tree growth in our study is consistent with expectations from stomatal conductance models (Grossiord *et al*. 2020), with stomatal closure and ensuing restriction of CO_2_ assimilation rate triggered by VPD values exceeding the climate mean and usual variation range. This negative effect of VPD is expected to be amplified by SRAD anomalies, as VPD depends on leaf temperature, which itself increases with SRAD (Grossiord *et al*. 2020). The negative influence of SRAD anomalies on tree growth may be additive to that of VPD anomalies, as previously shown (Rifai *et al*. 2018, 2019; Krause & Winter 2020). Furthermore, positive SRAD anomalies did not enhance tree growth but reduced it, as would be expected if its effect was VPD-related. However, the effect of SRAD anomalies on tree growth was probably more than a mere reflection of VPD, as anomalies in SRAD and VPD were only moderately correlated (*r* = 0.33, Table S3a). Excess or fluctuating light, and changes in light quantity and quality are other potential mechanisms underlying SRAD anomaly effects, as these can be direct physiological stressors (Krause & Winter 2020; Roeber *et al*. 2020), or indirectly influence the response to other abiotic or biotic stresses (Roeber *et al*. 2020).

The strong effect of VPD anomalies compared to the undetectable effect of MCWD anomalies suggests that VPD may limit tree growth before soil water becomes limiting, further confirming previous results in temperate and tropical forests (Choat *et al*. 2012; Novick *et al*. 2016; Konings *et al*. 2017; Rifai *et al*. 2018; Sanginés de Cárcer *et al*. 2018). This is a key result, given the generalised tree growth decrease potentially driven by increasing VPD anomalies, as VPD has been strongly increasing in the tropics due to anthropogenic climate change (Rifai *et al*. 2019). Yuan et al. (Yuan *et al*. 2019) highlighted a particularly-strong increasing VPD trend at the global scale beginning in the late 1990’s (0.017 hPa / yr). Modelling VPD anomalies through time from 2000 to 2019 in our dataset, we detected a 3.8-fold stronger VPD increase rate across all plots (0.045 hPa / yr, 90%-HPDI: 0.019, 0.066; R² = 0.80; details in Table S4; e.g. Fig. 1b). This trend itself was stronger than the 1971-2019 trend in our dataset (0.020 hPa / yr; R² = 0.84; Table S4), indicating a sharper-than-previously-thought VPD increase in the past two decades. This rapid increase of VPD anomalies through time combined with the generalised ensuing decrease in tree growth and growth sensitivity variability to VPD among species (Fig. S3; Table S5) suggests that tropical forest composition and functions may be strongly altered by ongoing climate change, especially by VPD. It is worth noting that soil water deficit also depends on evapotranspiration estimates accuracy and variables unaccounted for, here (e.g. soil water retention capacity, topography), so that the importance of soil-related water stresses should be interpreted with caution.

In spite of the suppressing effects of increasing anomalies in SRAD, VPD, and Tmean, average growth rates were higher in warmer and sunnier forests (i.e. higher long-term means), across species (Fig. 2) and within many species (Table S5). While long-term Tmean was highly correlated with elevation (*r* = -0.95; Table S3a), mean solar radiation was not correlated with neither elevation nor the other climate variables (Table S3a). This suggests that these forests are in general energy-limited along the elevation gradient (faster growth in lowland forests), and light-limited across the gradient, supporting previous results along an Amazon-Andes elevation gradient (Fyllas *et al*. 2017). Our gradient of mean climates encompassed 7 to 51% of the global-scale climate space of tropical forests, but did not encompass their driest and warmest conditions (see Fig. S7). Future studies will need to cover a broader range of climate values to test how generalisable the relationships that we detected are for tropical forests worldwide.

### Tradeoffs in tree growth responses to climate

We showed that two aspects allowed understanding the broad range of species differences in growth response to VPD anomalies: the long-term mean VPD where species grew better, and the contrast between slow- and fast-growing species (Fig. 3a, c). The models including plot-specific responses to climate anomalies additionally showed that forest growth sensitivity to VPD anomalies was stronger in drier forests, mostly at the higher end of the VPD range (Fig. 4a). This result could be driven by higher levels of obligate or facultative deciduousness, as even the wettest rainforests have seasonal peaks in leaffall (Edwards *et al*. 2018) and the drier the forest the earlier the leaffall peak and the shorter the growing season. Our results support recent findings indicating that drier forests could be more sensitive to increasing VPD anomalies (Aguirre-Gutiérrez *et al*. 2020; Powers *et al*. 2020), which would here translate into drier rainforests already being under water stress and therefore closer to a threshold of further growth decrease than moist rainforests. This effect may not be linear and will need to be further tested with more plots encompassing diverse water-stress conditions.

Similarly, Sullivan et al. (2020) recently showed that warmer forests may be closer to a temperature threshold beyond which woody productivity would decrease. In our study, this would translate into expectations that forests and species adapted to warmer conditions would respond more negatively to further temperature increases. Our results are consistent with this expectation but suggest that the temperature effect manifests itself indirectly through VPD.

Species that grew faster in cloudier forests showed the strongest growth reduction due to positive SRAD anomalies (Fig. 3b). This may reflect species differences in light-use strategies, with species that grow well under low direct-sunlight conditions not benefitting from brighter conditions. This was supported by the stronger negative effects of SRAD anomalies in species with lower maximum photosynthetic capacity, stomatal conductance and electron transport capacity (Fig. 5), a trait syndrome consistent with shade-tolerance strategies (He *et al*. 2019). This interpretation was supported in the plot-level analyses by the steeper growth rate decreases in the cloudier forests in response to positive SRAD anomalies (Fig. 4b), which may stem from a plot-wide relatively more marked adaptation to shade tolerance.

### Functional traits mediate the effects of climate anomalies on tree growth

Species traits directly influenced species intrinsic growth rate. As expected, intrinsic growth rate increased with metabolism (Rd), maximum size (DBH_max_), and acquisitive chemical and physiological traits related to the photosynthetic machinery (leaf P content, Asat and Vcmax). However, it also increased with leaf δ^13^C and LMA, contrary to expectations as high values of these traits correspond to tough, long-lived leaves and high intrinsic water use efficiency (Cernusak *et al*. 2013; Osnas *et al*. 2013). In our study, leaf δ^13^C was positively correlated with leaf N and P contents (Table S3b), suggesting variation in δ^13^C among species may have been driven more by photosynthetic capacity than by stomatal conductance. The positive association of LMA and growth, also reported in previous studies (Poorter *et al*. 2008; Wills *et al*. 2018; Gray *et al*. 2019), could be explained by a change in the cost-benefit balance of acquisitive traits with plant size (Gibert *et al*. 2016; Gray *et al*. 2019).

An overarching finding is that species traits can enhance our understanding of differences in species growth response to the anomalies of SRAD and VPD, and to forest stand structure. Our results confirmed that resource-acquisitive species overall had higher intrinsic growth rate and that their growth was more sensitive to positive anomalies in SRAD and VPD. This highlights a tradeoff between fast growth (via high allocation to acquisitive tissues) and sensitivity to atmospheric water stress, consistent with expectations from the ‘fast-slow’ plant economics spectrum (Reich 2014). Most physiological traits directly related to photosynthesis (Table 1) successfully captured species differences in growth sensitivity to SRAD anomalies (Fig. 5; Fig. S6), confirming the importance of physiological traits to investigate potential mechanisms underlying differences in demographic responses to climate change among species (Brodribb *et al*. 2020; Powers *et al*. 2020). Increasing values of these traits attenuated the tree growth reduction following increasing SRAD anomalies (Fig. 5; Fig. S7), suggesting that species investing in a more responsive and flexible photosynthetic machinery may cope better with unusually-high direct exposure to sunlight. While most traits that increased species intrinsic growth rate also exacerbated the negative effects of VPD anomalies on tree growth, the mediation of SRAD anomalies by species traits was mostly independent of the fast-slow spectrum (Fig. 5; Fig. S5, S6). For example, while leaf P concentration, stable carbon isotope ratio and the maximum photosynthetic capacity tended to increase intrinsic growth rate, the two former accentuated while the latter attenuated the negative effects of SRAD anomalies on tree growth (Fig. 5).

### Stand structure as driver of tree growth variation

Plot basal area consistently strongly reduced tree growth across species and explained more growth variation than mean climate for all four climate variables. Although plot basal area was partly correlated with elevation, the 30-year average of Tmean and VPD (*r* = 0.63, -0.59, and -0.47, respectively; Table S3a), the slope coefficient of basal area remained virtually unchanged across models including Tmean, VPD, or the other less correlated covariates (and was much steeper than the slopes of long-term Tmean or VPD), so that the stand structure effect detected here is unlikely to indirectly reflect Tmean or VPD. Furthermore, faster growth in less dense environments across forest plots suggests a release from competition for light. This is supported by the general light-limitation suggested by the faster growth in sunnier sites. Slower growth in denser environments may also suggest an increase in competition for resources or attacks by natural enemies. Neighbourhood crowding has indeed been shown to strongly reduce tree growth in tropical and temperate forests (Clark *et al*. 2014; Fortunel *et al*. 2016, 2018; Uriarte *et al*. 2016). In line with these studies, we found that conservative species with high wood density suffered less growth reduction from increasing plot basal area, while acquisitive species with high dark respiration rate and leaf δ^13^C were more sensitive to basal area (Fig. S5, S6).

In summary, we have shown how long-term demographic data across multiple plots encompassing environmental gradients, combined with functional traits collection can yield insights into how climate affects interannual variation of tree growth at different temporal scales, and give important clues into which species and forests may be particularly vulnerable to climate change, and why. Our findings emphasise the importance of functional traits - and notably those directly related to photosynthesis and water use efficiency - to understand species differences in demographic sensitivity to abiotic and biotic drivers. Future efforts to further characterise how climate and neighbourhood crowding affect tree growth, survival, and population growth across environmental gradients, and how these effects are mediated by species traits will help improve predictions of forest response and future ecosystem functions to climate change under different trajectories.

## Acknowledgments

We thank Alex Cheesman for his help with field work in Bellenden Ker. DB and GD were supported by the Wiener Anspach Foundation. YM was supported by the Frank Jackson Foundation. The trait campaign and data analysis were funded by UK Natural Environmental Research Council (NERC) Grant NE/P001092/1 (to YM) and European Research Council projects T-FORCES (Tropical Forests in the Changing Earth System) to YM and OLP. and GEM-TRAIT to YM. We are thankful to the Daintree Rainforest Observatory for providing a subsidy on accommodation and station fees.

## Statement of authorship

DB, CF and YM designed the study. DB tidied and vetted the demographic and trait data and performed the analyses. D.B. and C.F. designed the statistical models of tree growth. S.R. helped generating the climatic covariates and created Fig.1a. I.O. and JAG contributed ideas and constructive feedback to early versions of the work. JAG helped obtain climate data and provided feedback on an early version of the work. L.C. and L.P.B. led the trait data collection, assisted by RD, BEM., HN, JC and PS. MH provided the final raw climate data. MB supplied demographic data for the 20 CSIRO plots and Robson Creek. LC, SR, JAG, GD provided feedback to part of the discussion. SGWL contributed demographic data of Daintree Observatory. DB led the writing with regular feedback from YM, CF and SMM on intermediate stages of the analyses and manuscript. All authors commented on the manuscript and gave their approval for the publication.

## Data accessibility statement

The raw demographic data that supported the findings are available in Bradford et al. (2014) and CSIRO Data Access Portal (https://data.csiro.au/dap/). R code, raw and processed data will be archived in a Dryad repository, whose DOI will be added at the end of the article, and will be available from the corresponding author upon request.

## Supplementary Materials

Figure S1: Trait turnover along the elevation gradient.

Figure S2: Effects of long-term climate average, short-term anomalies, tree size and stand structure on tree growth rate.

Figure S3: Illustration of the variability of tree growth responses to the climatic drivers among species.

Figure S4 Coordination among the species-level growth responses to stand structure and intrinsic growth rate.

Figure S5: Effects of tree size, mean climate, climate anomalies, stand structure, and species functional traits on intrinsic growth rate.

Figure S6: Mediation of climatic and stand structure effects on tree growth by species functional traits.

Figure S7: Overlap of the climate spaces of the 23 studied tropical rainforest plots and tropical wet forests worldwide.

Supplementary Tables S1-S5: Forest plots, species characteristics, covariate correlations, and statistical model detailed outputs.

Supplementary Methods S1: Extended Material and Methods

Supplementary Methods S2: R code for the calculation of climate covariates and construction of the individual tree growth models.

## 1. Supplementary Figures

Fig. S1 shows how trait value distributions change across the elevation gradient. All photosynthetic traits (but Amax) tend to increase with elevation, and so do LMA, leaf thickness and wood density. Dark respiration rate, leaf δ^13^C, P_leaf_ and leaf area decrease with elevation. Although most traits see a significant increase or decrease in their values, the whole trait range remains well represented across the whole gradient. In addition, it is worth noting that tree growth decreased with elevation (Fig. 2; positive effect of historical mean Tmean on growth, which is strongly negatively correlated to elevation, *r* = -0.95, see Table S3a), while several traits that increased with elevation (Fig. S1) also had a positive effect on intrinsic growth rate (Fig. 5 and Fig. S5e) (e.g. Vcmax, LMA). This indicates that the effects of traits on growth are actual trait effects rather than indirect elevation effects. This is confirmed by the fact that the trait effects on intrinsic growth rate (from the model including Tmean as climate variable) (Fig. S5e) are also detected in the models including a different climate variable (and no proxy of elevation; see Fig. S6i-l).

**Figure S1:**
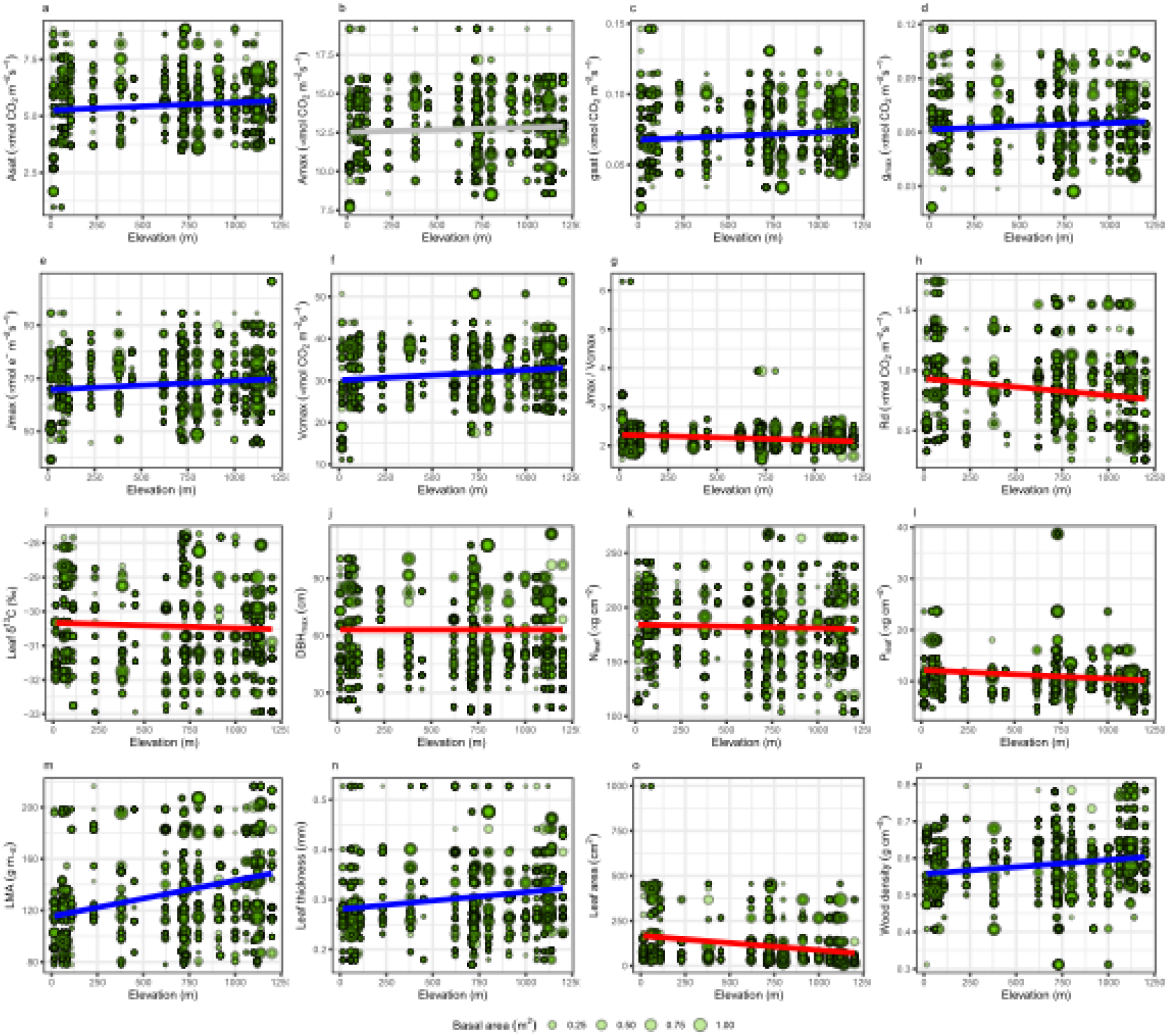
Trait turnover along the elevation gradient. Trait turnover along the elevation gradient. Circles are individual trees. Circle diameters are proportional to individuals’ average basal area across the multiple censuses. Regression lines were drawn from linear regressions of the traits against elevation, using a frequentist approach, where observations (circles) were weighted by their basal area. The significance threshold (*p*-value of 0.05) was adjusted for multiple tests using the Sidak correction. Red and blue lines are significant negative and positive elevation effects, respectively. Grey lines correspond to non-significant tests.

**Figure S2:**
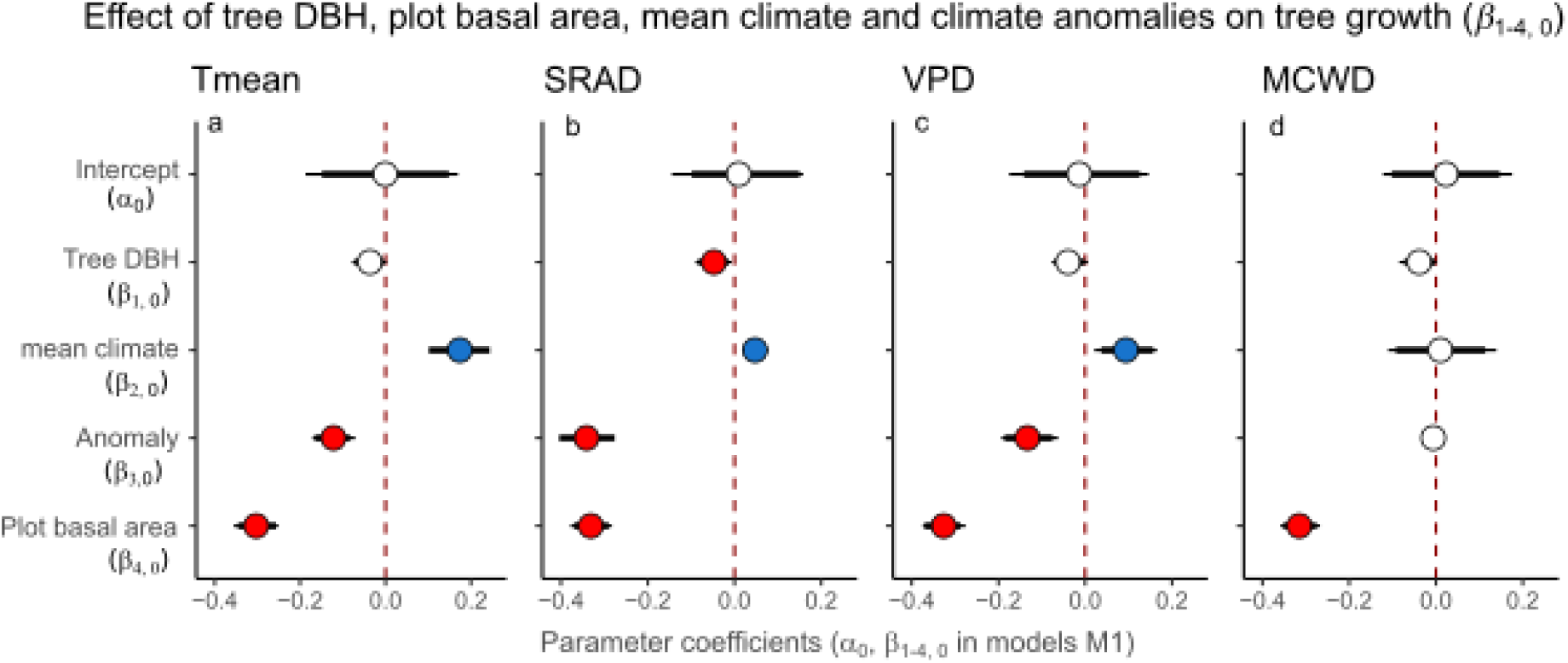
Effects of long-term climate mean, climate anomalies, tree size and stand structure on tree growth rate (models based on all 509 species; no trait) Grandl effects of climate, stand structure and tree DBH on tree growth in the separate models including Tmean (a), SRAD (b), VPD (c), and MCWD (d) (M1 models on all 509 species; no trait; see eqs. 3 for coefficient codes). Circles, thick and thin intervals are median, 90%- and 95%-HPDI of coefficient posterior probability distributions. Red and blue circles indicate negative and positive effects on tree growth, respectively, for the covariates with clear effects (95%-HPDI not encompassing zero); white circles indicate coefficients whose 95%-HPDI include zero.

**Figure S3:**
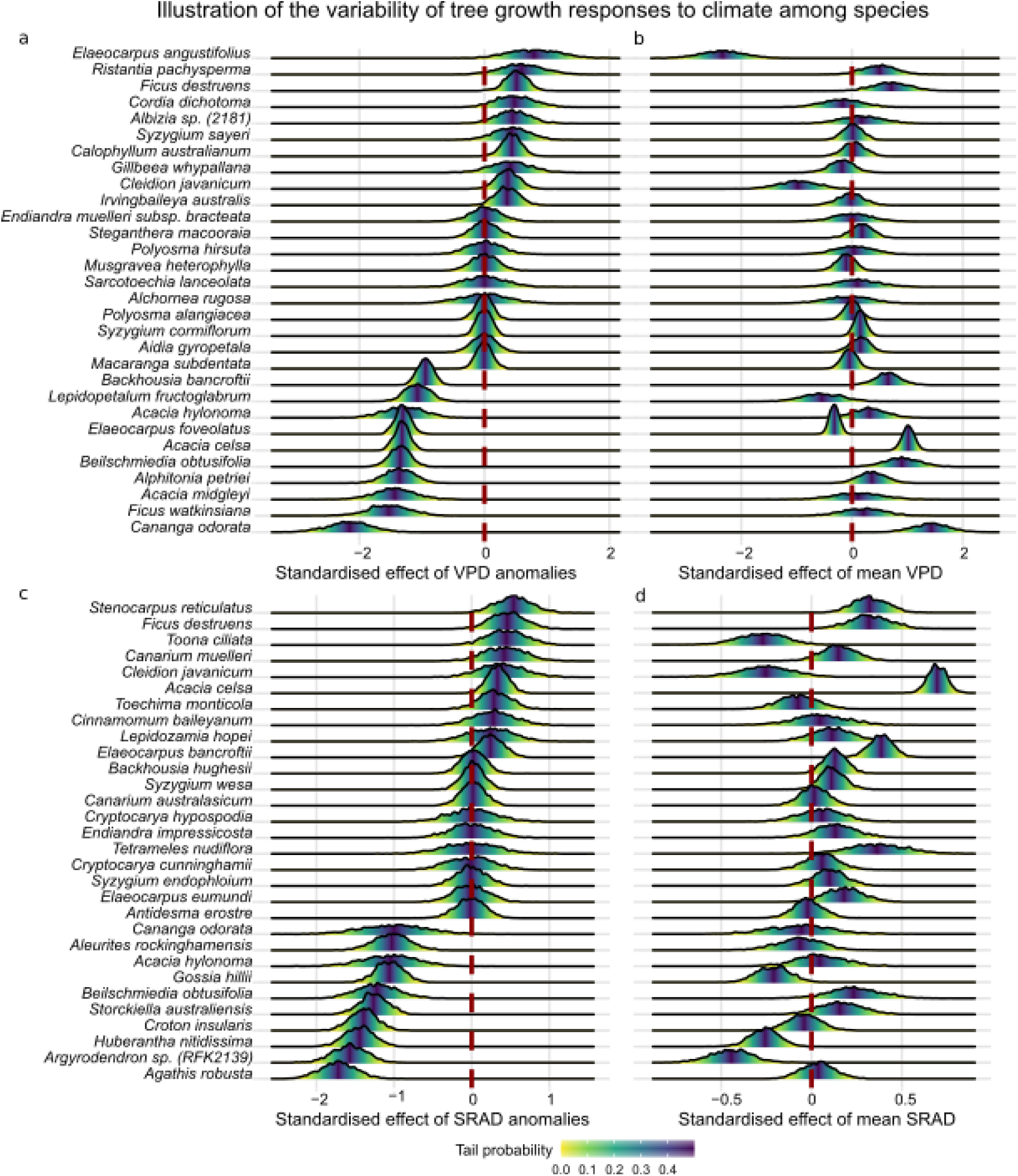
Illustration of the variability of tree growth responses to the climatic drivers among species. Illustration of interspecific variability in tree growth sensitivity to the climatic drivers. Fig. S3a-d illustrate the species-level posterior distributions of the slope coefficients associated to four model covariates for 30 species (among the 509). Fig. S3a,c: Tree growth response to VPD and solar radiation anomalies, displaying the 10 species presenting the strongest positive response, followed by 10 species presenting no particular response, and the 10 presenting the strongest negative response. Fig. S3b,d display the same 30 species for their tree growth response to the long-term mean VPD and solar radiation (1981-2010), respectively. Species-level growth responses to all the model covariates for all 509 species (i.e. species-level slope coefficients) are in Table S4. The vertical dashed lines separate positive responses (i.e. growth rate increase with the corresponding covariate) on the right from negative responses on the left. Species whose posterior distribution does not include zero at all, or include it in the yellow tail of the posterior, can be considered as responding clearly to the corresponding climate covariate (90% of the posterior probability mass of the slope value is smaller or higher than zero). Comparing the vertical ordering of the posteriors between a and b, and between c and d, shows part of the significant correlations between the species-level growth sensitivities to multiple drivers corresponding to Fig. 3a and Fig. 3b.

**Figure S4:**
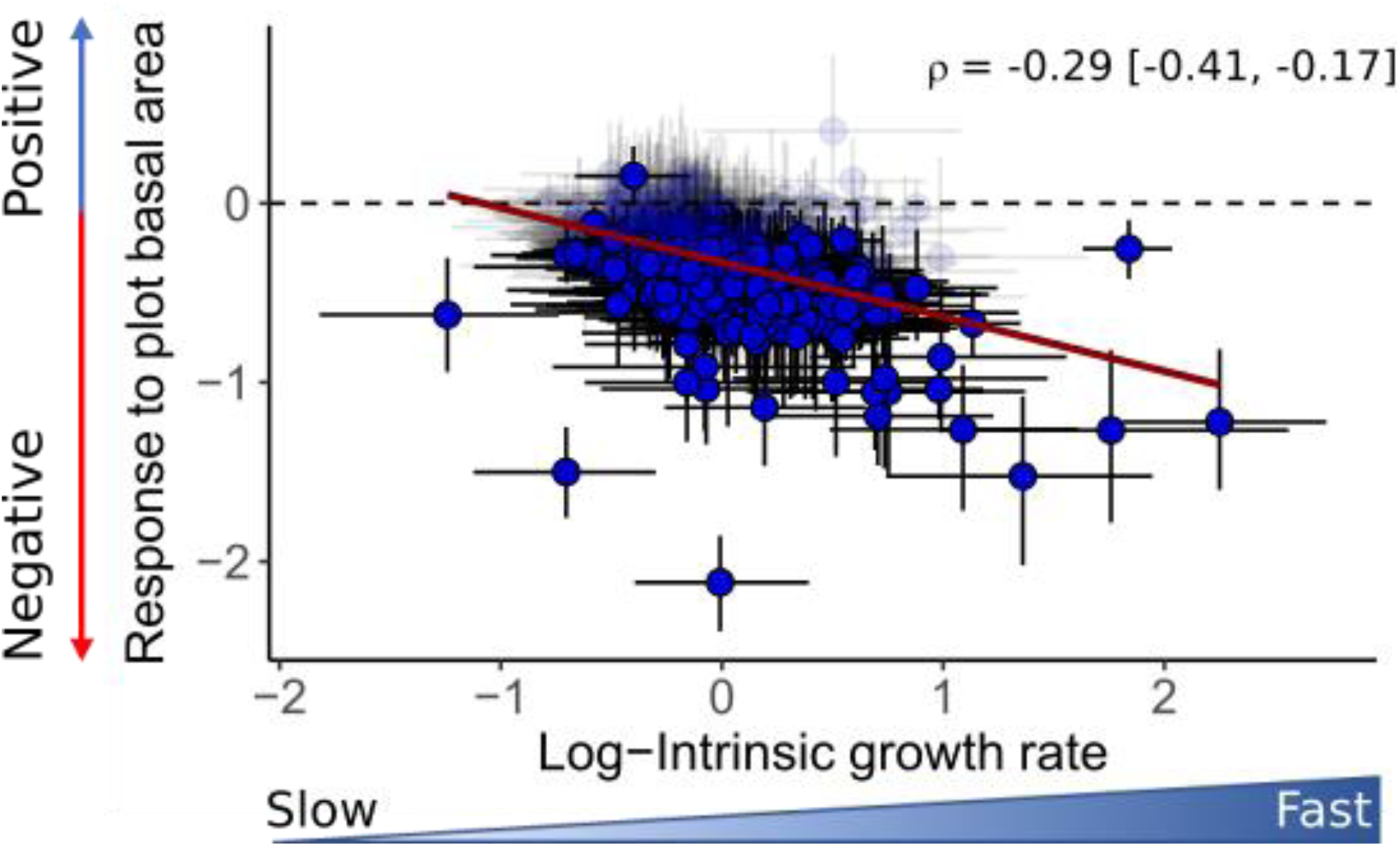
Coordination among the species-level growth responses to stand structure and intrinsic growth rate. Correlation among species-level growth responses highlighting joint responses to multiple drivers (models of all 509 species; *ρ*, see eq. 3.5 in Supplementary Methods S1). Species-level correlation between tree growth sensitivity to plot basal area and intrinsic growth rate. Circles are species, placed at the median of their corresponding coefficient posteriors. Vertical and horizontal bars are 95%-HPDI for the corresponding coefficients. Species for which both plotted coefficients were clearly different than zero (95%-HPDI not encompassing zero) are plain blue; other species are shaded. The red regression line indicates a clear negative correlation (95%-HPDI of the correlation posterior not encompassing zero; mean, lower and upper 95%-HPD interval values provided in the upper right-hand corner of the plots). On the *y*-axis, values above and below zero indicate positive and negative effects of plot basal area on growth, respectively.

Fig. S4 shows that tree growth is more reduced by high plot basal area among fast-growing species than among slow-growing species, suggesting a trade-off between a fast average growth rate but higher sensitivity to competition for light or other resources (or natural enemies), and less sensitivity to stand structure but a slower average growth rate (see *Discussion*).

**Figure S5:**
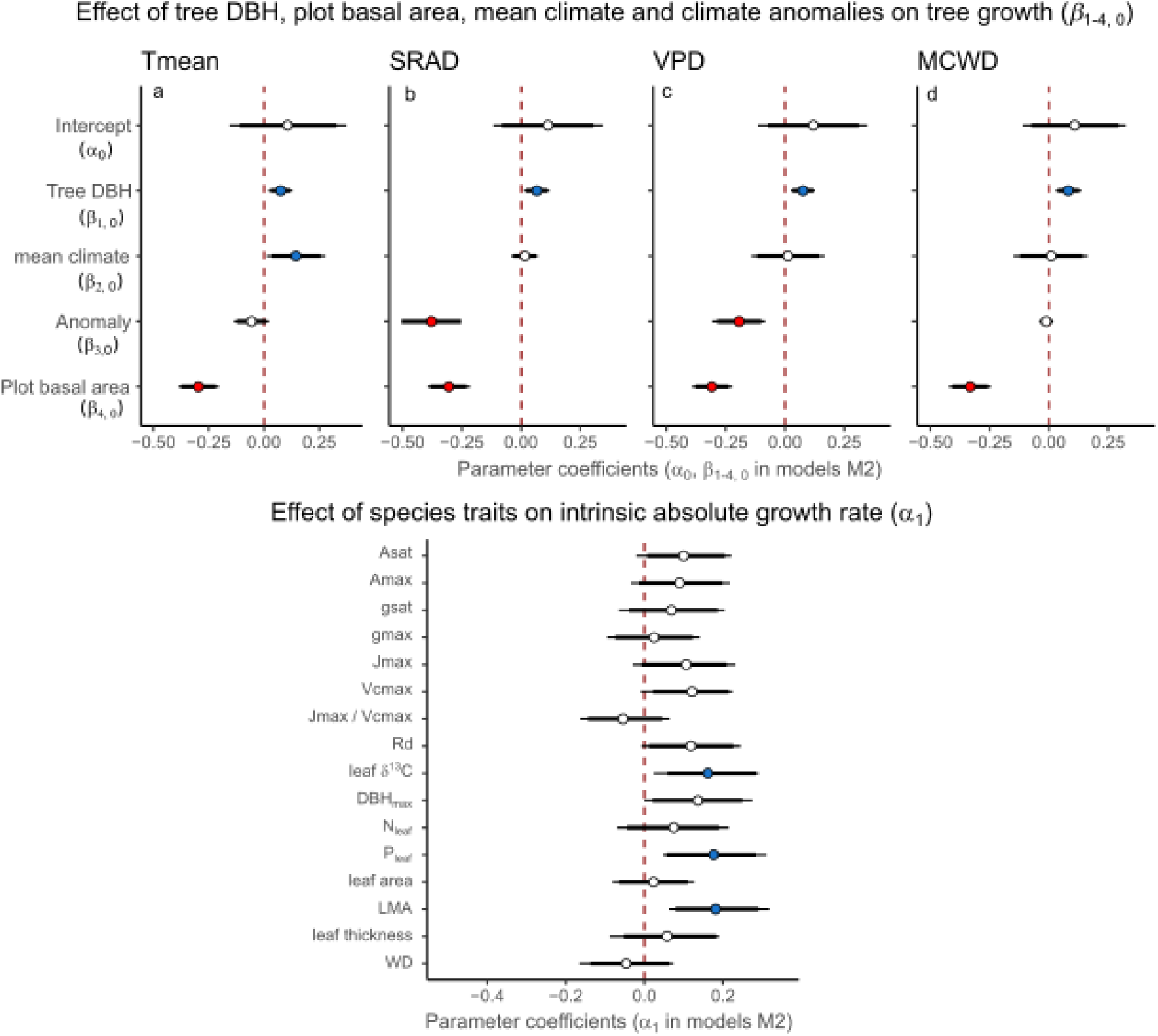
Effects of tree size, mean climate, climate anomalies, stand structure, and species functional traits on intrinsic growth rate (based on the 75 species with measured trait data) Influences of tree size, plot basal area, mean climate and climate anomalies (a-d; same as Fig. S2b-e) and effects of species traits on intrinsic tree growth (e; M2 models, using 75 species with measured trait data; see coefficients codes in eqs. 4, Supplementary Methods S1). Models were run separately for mean temperature (Tmean), vapour pressure deficit (VPD), maximum climatological water deficit (MCWD) and solar radiation (SRAD), each model containing the climate mean (1981-2010) and anomalies of the corresponding climate variable (see *Methods*). Each of these four models were run with each trait separately. The indirect effects of traits on tree growth through their mediation of climate and stand structure effects are not represented here, for clarity, but are in Fig. S6. Circles, thick and thin intervals are median, 90%- and 95%-HPDI of coefficient posterior probability distributions. Red and blue circles indicate negative and positive effects on tree growth, respectively, for the covariates with clear effects (95%-HPDI not encompassing zero); white circles indicate coefficients whose 95%-HPDI include zero. e: Trait effects on intrinsic growth rate (i.e. on the intercept; α_1_ coefficient) from the model including VPD (Fig. S5c). The direct traits effects on growth rate from other models (a, b, d) were similar and not shown here for clarity.

The models run on the 75 species for which trait data were measured yielded similar results than the models run on all 509 species and without trait effects, regarding the climatic and stand structure covariate effects, with some differences. The effect of tree size, negative overall based on all 509 species (Fig. 2), became positive. This indicates that while tree growth rate still decreased with tree size in some species (Table S5), it increased with tree size for a large proportion of the 75 species. While showing the same trends, the negative effect of the anomalies in temperature became unimportant (95%-HPDI encompasses zero), and so did the positive effect of mean VPD and solar radiation.

Species intrinsic growth rate increased with leaf δ^13^C, LMA, leaf P content (95%-HPDI not encompassing zero), and dark respiration rate (Rd), DBH_max_, Asat and Vcmax (90%-HPDI not encompassing zero).

**Figure S6:**
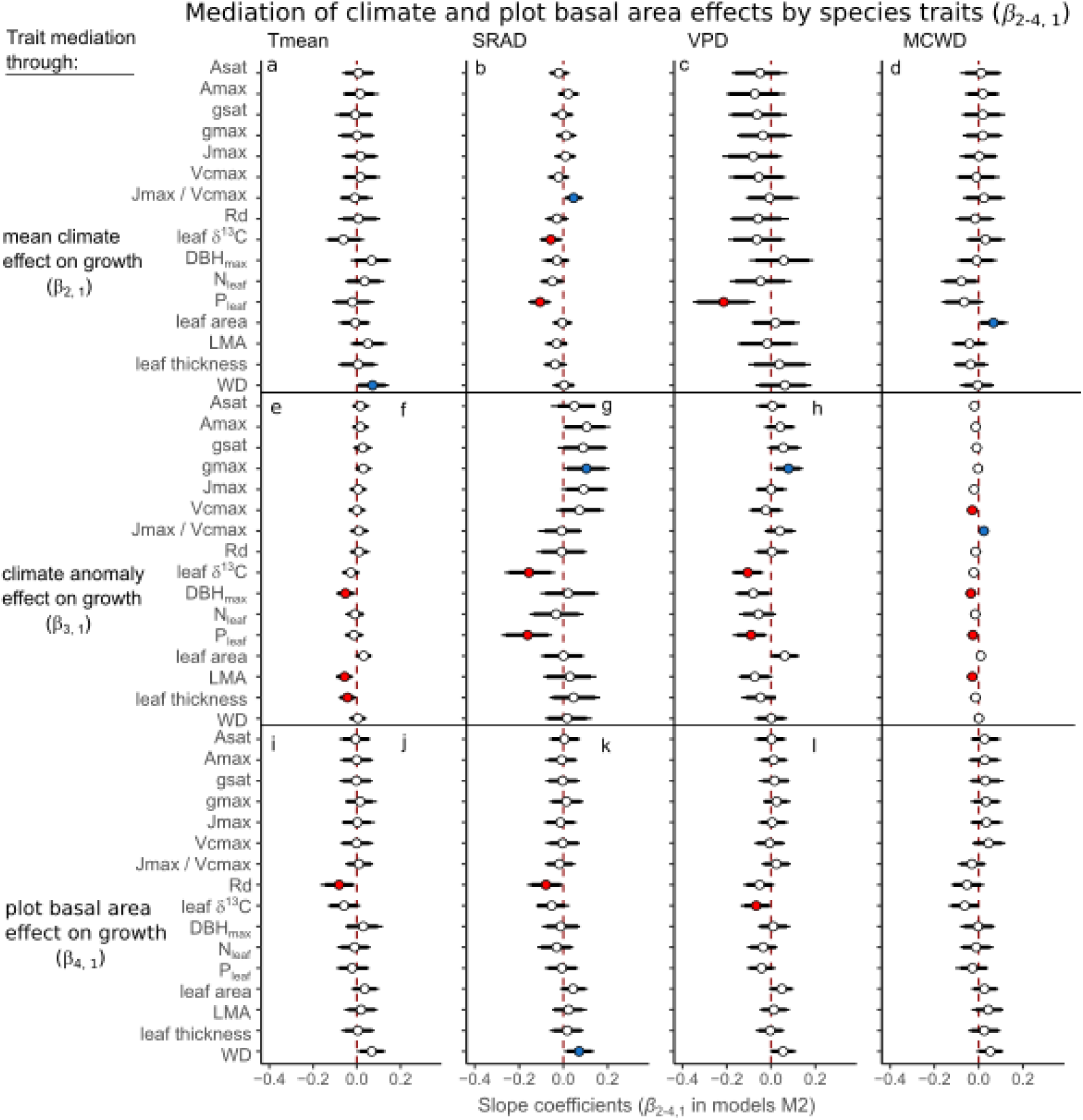
Mediation of climatic and stand structure effects on tree growth by species functional traits (M2 models, from 75 species with trait data) Effects of species traits on the influences that mean climate (β_2,1_), climate anomalies (β_3,1_) and plot basal area (β_4,1_) have on tree growth (based on the 75 trait species; see eqs. 4 in Supplementary Methods S1). Models were run separately for Tmean, VPD, MCWD and SRAD. Models were run with each trait separately. Circles, thick and thin intervals are median, 90%- and 95%-HPDI of coefficient posterior probability distributions. Red and blue circles indicate negative and positive effects on tree growth, respectively, for the covariates with clear effects (95%-HPDI not encompassing zero); white circles indicate coefficients whose 95%-HPDI include zero. Refer to climate and stand structure effects (Fig. S5) to define whether traits accentuate (trait mediation effect on Fig. S6 has the same sign than the direct covariate effect on Fig. S5) or attenuate (opposite sign) the effects of the main drivers on growth (see *Methods* for details).

**Figure S7:**
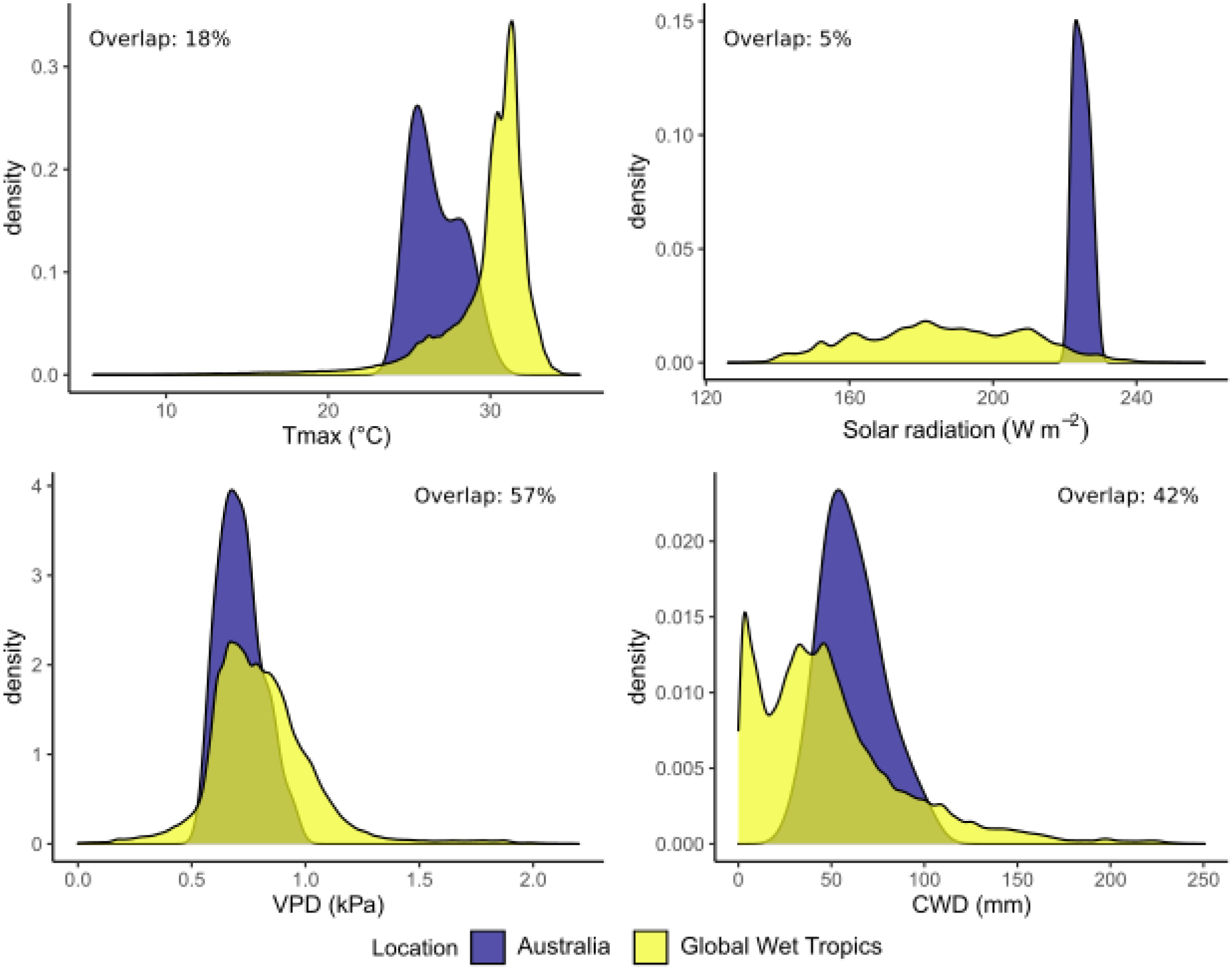
Overlap of the climate spaces of the 23 studied tropical rainforest plots and tropical forests worldwide. Comparison of the climatic space occupied by the 23 permanent plots of tropical rainforests of the study with the total climatic space of tropical wet forests worldwide. The climatic spaces were obtained from 30-year climate averages (1985-2015) extracted from TerraClimate (Abatzoglou *et al*. 2018), combined with the spatial locations of the grid cells belonging to the ecoregion “Tropical and subtropical moist broadleaf forests” ((Dinerstein *et al*. 2017); see https://ecoregions2017.appspot.com/).

## 2. Supplementary Tables

The supplementary Table S1-S7 are in a separate Excel document.

## 3. Supplementary Methods S1

### Study sites and demographic data

Individual tree annual absolute growth rates were calculated for 12,853 trees in 23 permanent forest plots of tropical rainforest located in northern Queensland, Australia, between 12°44’ S to 21°15’ S and 143°15’ E to 148°33’ E, and encompassing an elevation gradient between 15 and 1200 m a.s.l. (Fig. 1a). Twenty of these plots (0.5-ha, 100 × 50 m) were established between 1971 and 1980 to provide long-term ecological and demographic data (Bradford *et al*. 2014), while three plots were established more recently along the same elevation gradient (Table S1). With two exceptions, all CSIRO permanent plots were established in unlogged forest; at establishment, EP9 and EP38 showed evidence of slight disturbance in a section of the respective plot due to selective logging at least 20 years prior (Bradford *et al*. 2014). Regular cyclonic disturbance contributes to the dynamics of the forests (Murphy *et al*. 2013). These forests cover a wide range of mean annual temperatures (19°C to 26.1°C), precipitations (1213 to 3563 mm), solar radiation (17.8 to 19.4 Mj m^-2^ day^-1^) and vapour pressure deficit (6.5 to 11.8 hPa) (Table S1). At plot establishment, all trees with stems ≥ 10 cm diameter at breast height (DBH) were mapped, identified to species level and measured for diameter. The 20 long-term plots were re-measured every two years for ten years, and then at three- to four-year intervals, with diameter, recruits and deaths recorded, summing up to 10 to 16 censuses per plot. The remaining four plots were established more recently, between 2001 and 2012, and were resampled one to three times (Table S1).

All available censuses were used to calculate individual annualised absolute growth rate (AGR) based on DBH at times 1 and 2 (dates; *t*_1_ and *t*_2_), as:

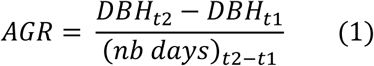

Abnormal AGR values were removed prior to analyses following (Condit *et al*. 2004). To do so, we removed the negative AGR values for which DBH*_t2_* was over four times SD_1_ below DBH*_t1_*, where SD_1_ = 0.0062 * DBH*_t2_* + 0.904 (in mm). These discarded values correspond to remeasurement of the wrong tree or to a digit dropped when encoding the data. The same correction could not be applied to positive AGR values, (see Condit *et al*. 2004 for details), so that we defined an upper AGR threshold value beyond which AGR values were considered outliers and removed.

*Syzygium graveolens* and *Elaeocarpus angustifolius* had a 95^th^ AGR percentile of 3.97 cm year^-1^ and 3.6 cm year^-1^, respectively. These were consistently the fastest growing species of the plot network. A total of 10 species presented and 95^th^ AGR percentile > 1.5 cm year^-1^. The threshold value for positive AGR outliers was set at 4 cm year^-1^.

### Climate data

We used four complementary climate variables relevant to tree growth and showing variability among the studied plots to investigate the effect of mean climate and climate anomalies on tree growth: mean temperature (Tmean), solar radiation (SRAD), vapour pressure deficit (VPD), and maximum climatological water deficit (MCWD).

Air evaporative demand – captured through VPD, the difference between air water vapour pressure at saturation and the actual water vapour pressure at a given temperature – can lead to reduced stomatal conductance while increasing evapotranspiration in many species, and can therefore affect photosynthesis and growth (Grossiord *et al*. 2020). Soil water deficit also controls tree growth through the balance between evapotranspiration and soil water availability, itself related to soil type and precipitation regime (Malhi *et al*. 2009). The MCWD is a proxy of the annual accumulated water stress over the drier season and is estimated from climate data as the cumulative deficit between precipitation and evapotranspiration, hence better capturing the seasonality of precipitation and potential soil water deficit than precipitation itself (Aragão *et al*. 2007; Malhi *et al*. 2009, 2015). Temperature partly controls photosynthesis, and increased temperatures can push species beyond their optimal conditions (Doughty & Goulden 2009; Brodribb *et al*. 2020), increase respiration costs (Tjoelker *et al*. 2001) and therefore change the proportion of photosynthates allocated to growth. Finally, solar radiation is directly related to the photosynthetic assimilation of CO_2_, so that increasing solar radiation could enhance tree growth through higher photosynthetic rates in these tropical rainforests (Fyllas *et al*. 2017), but could also reduce growth by indirectly increasing the leaf-to-air VPD (Grossiord *et al*. 2020).

Monthly climatic variables were obtained for the period 1970 to 2018 for each plot from ANUClimate v.2.0 (Hutchinson *et al*. 2014) (except for actual evapotranspiration), a spatial model constructed from a new anomaly-based approach to the interpolation of Australia’s national point climate data to produce climate variables on a 0.01° longitude-latitude grid. The monthly background means and the monthly anomaly values were spatially interpolated by trivariate thin plate smoothing spline functions of longitude, latitude and vertically exaggerated elevation using ANUSPLIN Version 4.6 (Hutchinson *et al*. 2014), using additional dependencies on proximity to the coast for the temperature and vapour pressure variables. Station elevations for the gridded min and max temperature (Tmin, Tmax), solar radiation (SRAD) and vapour pressure deficit (VPD) were obtained from local averages of 0.01° grid values from the GEODATA 9 second DEM version 3. Mean monthly temperature (Tmean) was obtained from Tmin and Tmax. Station elevations for the gridded rainfall (precip) were obtained from local averages of 0.05° grid values from the GEODATA 9 second DEM version 3 (Hutchinson *et al*. 2008). The VPD we used was an average of daily VPD at 9am and 3pm. The monthly actual evapotranspiration (aet) was derived for the same time period from TerraClimate (Abatzoglou *et al*. 2018), a gridded climate product that statistically downscales (ca. 4 km) a combination of the CRU TSv4.01 empirical climate interpolation and the JRA-55 climate reanalysis product. The aet was used in combination with precip to calculate the monthly climatological water deficit (CWD), a simple proxy of meteorologically-induced cumulative water stress (soil water deficit). The CWD was reset to zero at the wettest month of the year (maximum precip calculated from the plot climatology) and had an upper bound at 1000 mm. To calculate CWD, we used the plot-specific monthly aet historical mean (1981-2010 climatology) instead of the actual monthly aet estimations to avoid potential biases related to acclimation of trees to warmer and drier conditions across the long time span of the study (this only minimally changed CWD values). The CWD was used to calculate the maximum climatological water deficit of the year (MCWD), a measure of the peak dry season water deficit (Aragão *et al*. 2007; Malhi *et al*. 2009, 2015; Rifai *et al*. 2019). The MCWD was calculated on a monthly basis through a rolling maximum over the previous 12 months. Absolute values of MCWD were used to ease interpretations of its effect on AGR, so that the higher the MCWD the stronger the soil water deficit.

For each main variable (Tmean, VPD, MCWD, SRAD) in each forest plot, a monthly 30-year mean and standard deviation were calculated (1981-2010 period) (Table S1). On this basis, we calculated in each plot the monthly anomalies for each variable (i.e. monthly 30-year mean *μ* subtracted from monthly value) and divided them by their location-specific 30-year monthly standard deviation *σ*, yielding standardised anomalies (Aragão *et al*. 2007; Rifai *et al*. 2018):

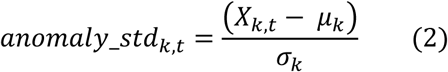

where X*_k, t_* is the climate variable value in plot *k* at time *t* (i.e. year and month), and *μ_k_* and *σ_k_* are the monthly 30-year mean and standard deviation of the corresponding plot’s location.

Standardised anomalies are expressed in units of standard deviation from monthly means over 1981-2010. This allows the comparison of plots differing not only in their historical means but also in the long-term variation range around them, that is, an important element to detect anomaly effects on tree growth across different climates. Note that calculating anomalies per month on the basis of the corresponding month 30-year mean ensures no possible confounding between anomalies and seasonal variability.

For the tree growth models, the monthly 30-year mean and standardised anomaly variables were averaged over the growth period between consecutive censuses (two to five years). For MCWD, the maximum over the growth period between two censuses was used instead of the weighted mean. The eight resulting interannual averaged variables were used as predictors to model tree growth (see *Data analysis*).

We did not include elevation in the growth models as it was already strongly correlated with long-term Tmean (*r* = -0.95; see Table S3a for correlations among climate variables, stand structure and elevation). Elevation was also somewhat correlated with VPD (−0.66), but was neither correlated with long-term solar radiation, MCWD, nor any of the anomaly variables (Table S3a). Among the standardised anomaly variables, Tmean and VPD were moderately correlated (*r* = 0.6), and smaller correlations were present between VPD and solar radiations (*r* = 0.32), and VPD and MCWD (*r* = 0.37). The chosen climate variables were therefore highly complementary and, besides long-term Tmean and elevation, were little correlated to one another or to the elevation gradient.

### Functional traits

Between July and September 2015, we measured the traits of 75 dominant, canopy trees in six of the 23 plots and two additional plots across the elevation gradient (Table 1; Table S1 and S2 for plot and species details, respectively). For each plot, species were chosen with the aim of sampling those that made up 80% of the standing biomass for the most recent census. Three individual trees were selected for each of these species. One sunlit branch was retrieved from the upper half of the crown from each of these trees by climbing and then using a pruning pole to excise the branch. Branches and leaves were chosen with minimal damage from herbivory. The branch was carried to a central measuring station, the cut end was submerged in a bucket of water, and then recut under water to remove any emboli introduced by the initial excision from the canopy. The cut surfaces of the branches remained submerged in the bucket throughout the course of the gas exchange measurements.

Five leaves, or leaflets in the case of compound leaves, were selected for gas exchange measurements with an LI-6400 portable photosynthesis system (Li-Cor Inc, Lincoln, NE, USA). Photosynthesis and stomatal conductance were first measured at a reference CO_2_ concentration of 400 µmol mol-1 and irradiance of 1500 µmol photons m^-2^ s^-1^ (*Asat* and *gsat*, respectively) supplied with an artificial light source (6400-02B LED, Li-Cor Inc.). A fixed block temperature was selected for each plot such that it was similar to the average daytime temperature at the time the plot was visited (19 to 30°C). Leaf-to-air vapour pressure difference during measurements was 1.1 ± 0.3 kPa (mean ± 1 SD). The CO_2_-saturated photosynthesis and stomatal conductance (*Amax* and *gmax*, respectively) were then measured at 1200 µmol mol^-1^ CO_2_. These measurements were repeated on the other four selected leaves or leaflets, and the resulting gas exchange parameters averaged for each branch.

One leaf per branch was wrapped in aluminium foil and left to dark-adapt for approximately 30 minutes, after which dark respiration (*Rd*) was measured. Block temperature was fixed as for the photosynthesis measurements. Leaf temperatures during dark respiration measurements ranged from 19 to 29°C. One leaf per species per plot was selected for measurement of a CO_2_ response (*A-c_i_*) curve. Temperature, irradiance, and leaf-to-air vapour pressure difference were as described, and the reference CO_2_ concentration was varied in the following sequence: 400, 250, 100, 50, 300, 400, 600, 800, 1200, and 1600 µmol mol^-1^, requiring two minutes for each step.

After gas exchange measurement, the leaves were scanned (Canon Lide 120) and leaf area was measured using Image J software (U. S. National Institutes of Health, Bethesda, Maryland, USA). Leaves were then dried at 70°C for 48 hours and their dry mass determined with an analytical balance (A&D Australasia, ANDW 464), for calculations of leaf mass per area. Leaf thickness was measured with a micrometer. A section of branch with diameter approximately 1 cm was then removed for wood density determination. Bark was removed and fresh volume determined by the water displacement method on an analytical balance (A&D Australasia, ANDW 464). The wood section was then dried and dry mass determined as for the leaves. In addition to the scanned leaves, approximately ten additional leaves were also collected from each branch and dried for determination of nutrient concentrations and stable isotope composition.

Dried leaf samples for each branch were bulked (without petioles) and ground to a fine powder using a laboratory mill (Cyclotec 1093, FOSS, Eden Prarie, MN, USA). They were analysed for concentrations of Ca, Mg, Na, K, B, Cu, Mn, Fe, Zn, P, and S by inductively coupled plasma optical emission spectrometry following peroxide assisted nitric acid digestion at a commercial laboratory (Nutrient Advantage, Werribee, Victoria, Australia). A separate aliquot from each branch was measured for stable carbon isotope ratio (δ^13^C), and total carbon and nitrogen concentrations, with an elemental analyser (CE Instruments, Milan, Italy) coupled to an isotope ratio mass spectrometer (Delta V; Thermo Fisher Scientific, Bremen, Germany) at the Advanced Analytical Laboratory, James Cook University, Cairns. The δ^13^C was expressed relative to the PeeDee Belemnite international standard.

The photosynthesis model of Farquhar et al. (Farquhar *et al*. 1980) was fitted to the *A*-c_i_ curves using the ‘plantecophys’ package in R (Duursma 2015), with estimates of the maximum carboxylation rate (Vcmax) and maximum light-driven electron flux (Jmax) normalized to 25°C. The one-point method (De Kauwe *et al*. 2016) was used to estimate Vcmax from net photosynthesis measured at 400 µmol mol^-1^ CO_2_, and Jmax from net photosynthesis measured at 1200 µmol mol^-1^ CO_2_ (Bloomfield *et al*. 2018). These estimates compared favourably to estimates from the full *A-c_i_* curves for the subset of branches on which both sets of measurements were conducted. The one-point estimates were therefore used in order to have estimates of these photosynthetic parameters for the full traits dataset.

### Data analysis

We addressed our questions through three sets of Bayesian multilevel models (M1 to M3).

#### M1: Tree growth response to climate means and anomalies, and species differences in their sensitivities to climate

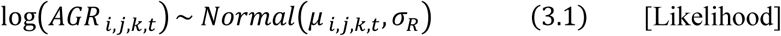

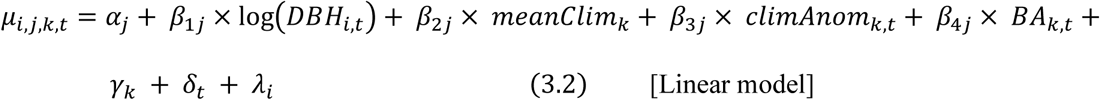

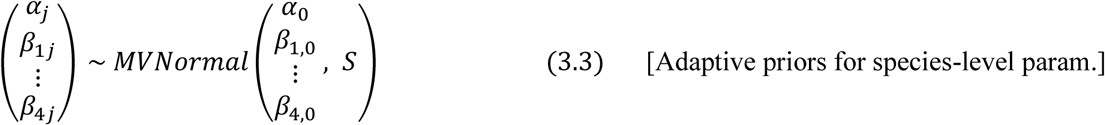

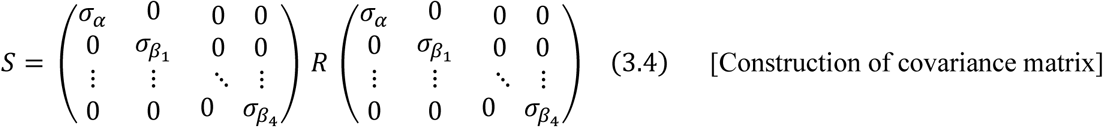

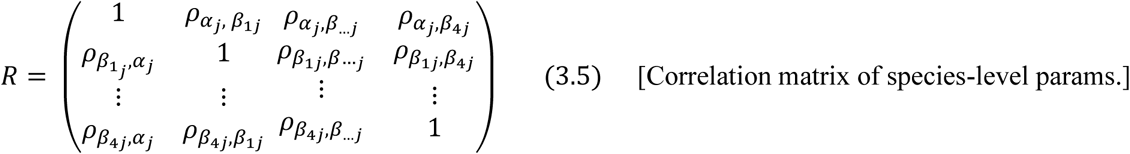

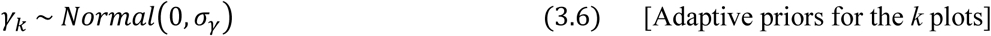

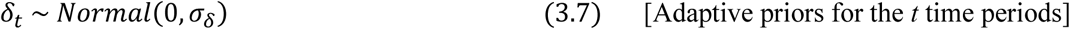

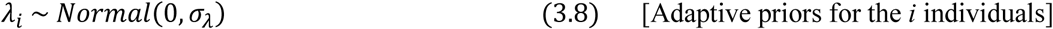

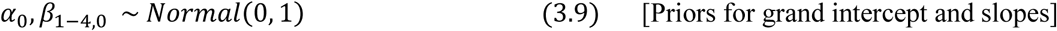

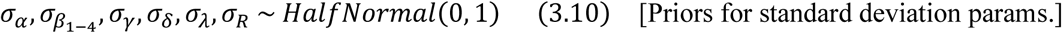

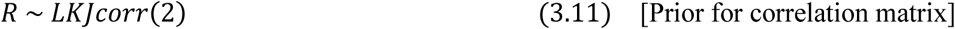

#### M2: Trait-mediated species-level tree growth response to climate

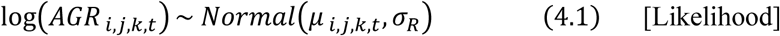

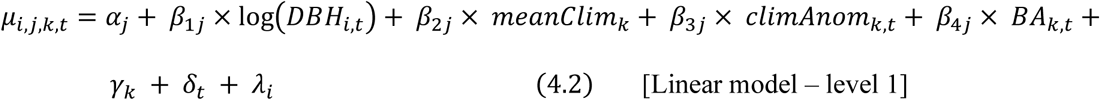

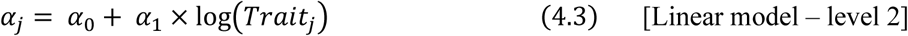

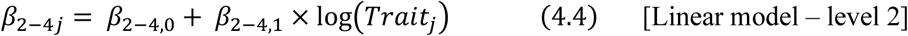

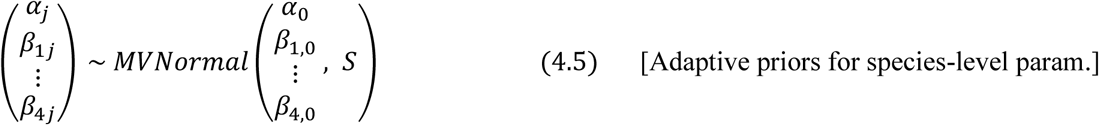

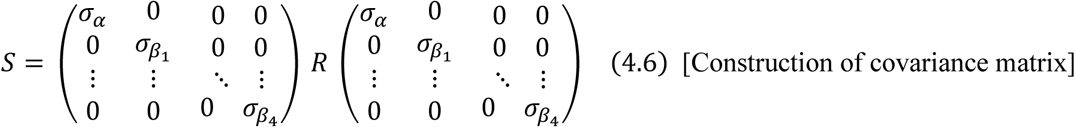

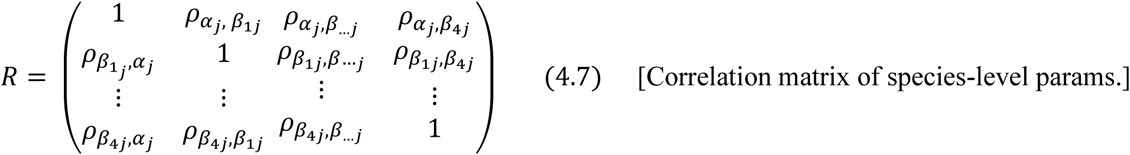

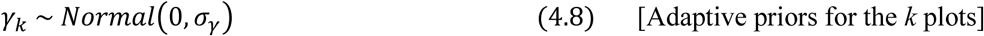

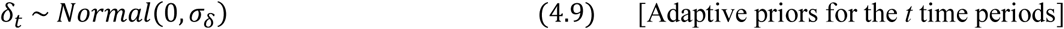

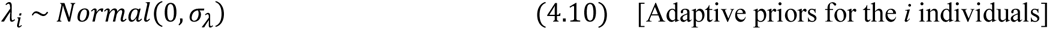

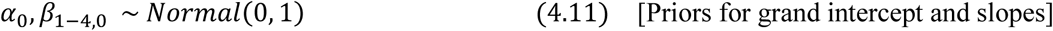

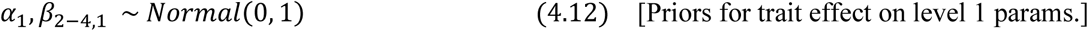

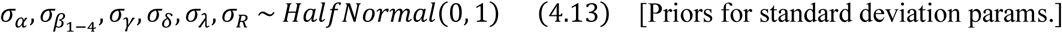

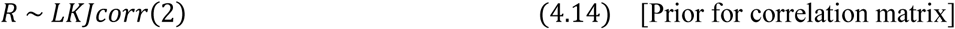

where eqs. 4.1, 4.2, 4.5-4.7 are the same as eqs. 3.1-3.5 of M1, while species-level intercepts and slopes are mediated by species mean trait value (eqs. 4.3-4.4; detailed equations and priors in Supplementary Methods S1). Parameter *α_1_* is the species-level departure from the grand intercept (*α_0_*) for an increase of one standard deviation in the log(trait T*_j_*) value of species *j* (direct effect of trait on AGR), while *β*_2-4, *1*_ are the departures from the grand slope of the corresponding model covariates for an increase of one standard deviation in the log(trait T*_j_*) value of species *j* (trait mediation of AGR response to climate and stand structure; see Supplementary Methods S1 for ecological interpretations of trait coefficient signs). We did not include the role of species traits in AGR response to tree size because some traits can change through tree ontogeny (Fortunel *et al*. 2020) and our trait data does not encompass species tree size ranges. M2 models were run separately for each of the four climate variables and for each of the 15 functional traits to manage model complexity (representing a total of 60 M2 models). For the covariates c ∈ (1, 2, 3, 4), negative and positive values of the *β*_c*j*_ or *β*_c,*0*_ slope parameters respectively indicated a negative and positive effect of the corresponding covariate on tree growth of species *j* (*β_cj_*) or across all species (*β_c,0_*). For instance, a negative *β*_3*,0*_ in the model including VPD as the climate variable would indicate that tree growth decreases when VPD anomalies increase, at the population level. If the sign of the trait coefficient (*β*_c, *1*_) is the same than that of the covariate it influenced (*β*_c, *j*_), then increasing values of trait T*_j_* accentuate the effect of covariate c on tree growth (i.e. push *β*_c*j*_ further away from 0). If the signs are different, increasing values of trait T*_j_* attenuate the effect of covariate c (i.e. pull *β*_c, *j*_ closer to 0).

In both M1 and M2 models, we standardised the response variable log(AGR*_i_*_, *j*, *k*, *t*_) and all covariates – but climate anomalies – to mean zero and unit standard deviation, to allow relative importance comparisons between covariates through slope coefficients (Schielzeth 2010), and to ease the assignment of plausible priors to the parameters (McElreath 2020) (eqs. 4.7-4.9). We did not standardise the averaged monthly anomalies to maintain their interpretability as deviations from long-term means in terms of plot-specific units of standard deviation (see eq. 2; i.e. mean anomaly covariate slope coefficients are not directly comparable to other covariate mean slopes). Individual trait measurements were averaged per species and log-transformed prior to standardisation, thus implying that parameter *β_c_*_, *j*_ corresponds to *β_c_*_, *0*_ at the mean trait value of the dataset.

#### M3: Plot-level tree growth response to climate anomalies and interaction with mean climate

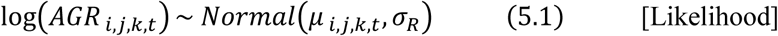

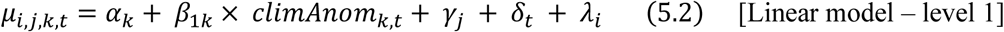

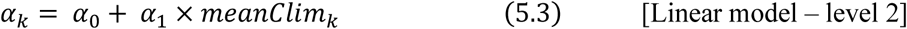

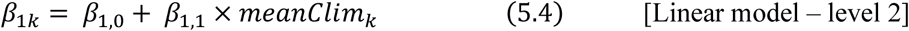

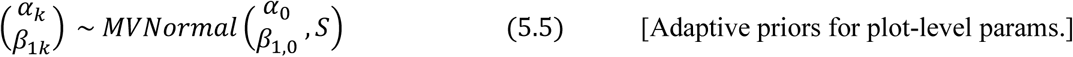

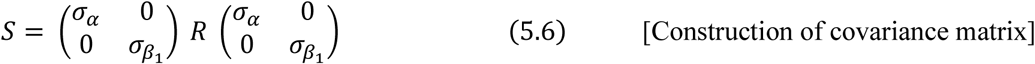

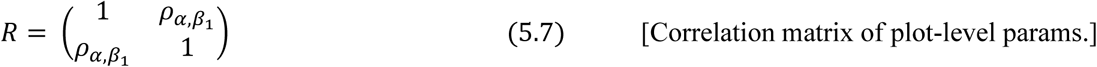

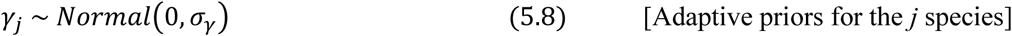

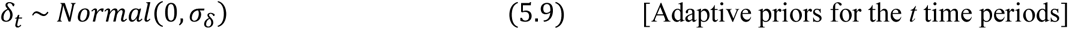

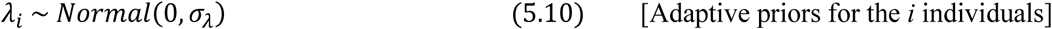

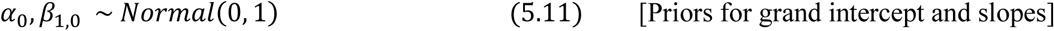

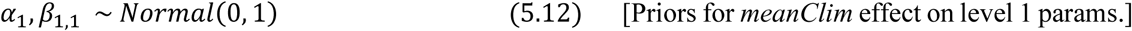

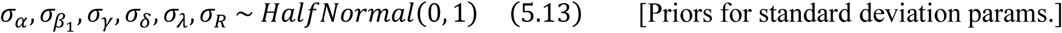

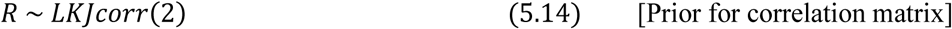

where *α_k_* is the average growth rate in plot *k*, and *β_1k_* characterises the growth response of plot *k* to standardised climate anomalies for time interval *t*. *α_0_* is the mean intercept value (i.e. mean absolute growth rate) across plots, and *α_1_* is the departure from the grand mean for one unit increase in mean climate. *β_1,0_* is the grand slope of climate anomalies, and *β_1,1_* is the departure from this grand mean for a one unit increase in mean climate (mediation of the effect of anomalies on growth by the plot mean climate; i.e., crosslevel interaction between the plot-level climate anomaly effect and the population-level mean climate effect). Parameters *γ_j_*, *δ_t_*, *λ_i_* are varying intercepts for species, census periods, and individual stems, respectively.

## 4. Supplementary Methods S2: R code

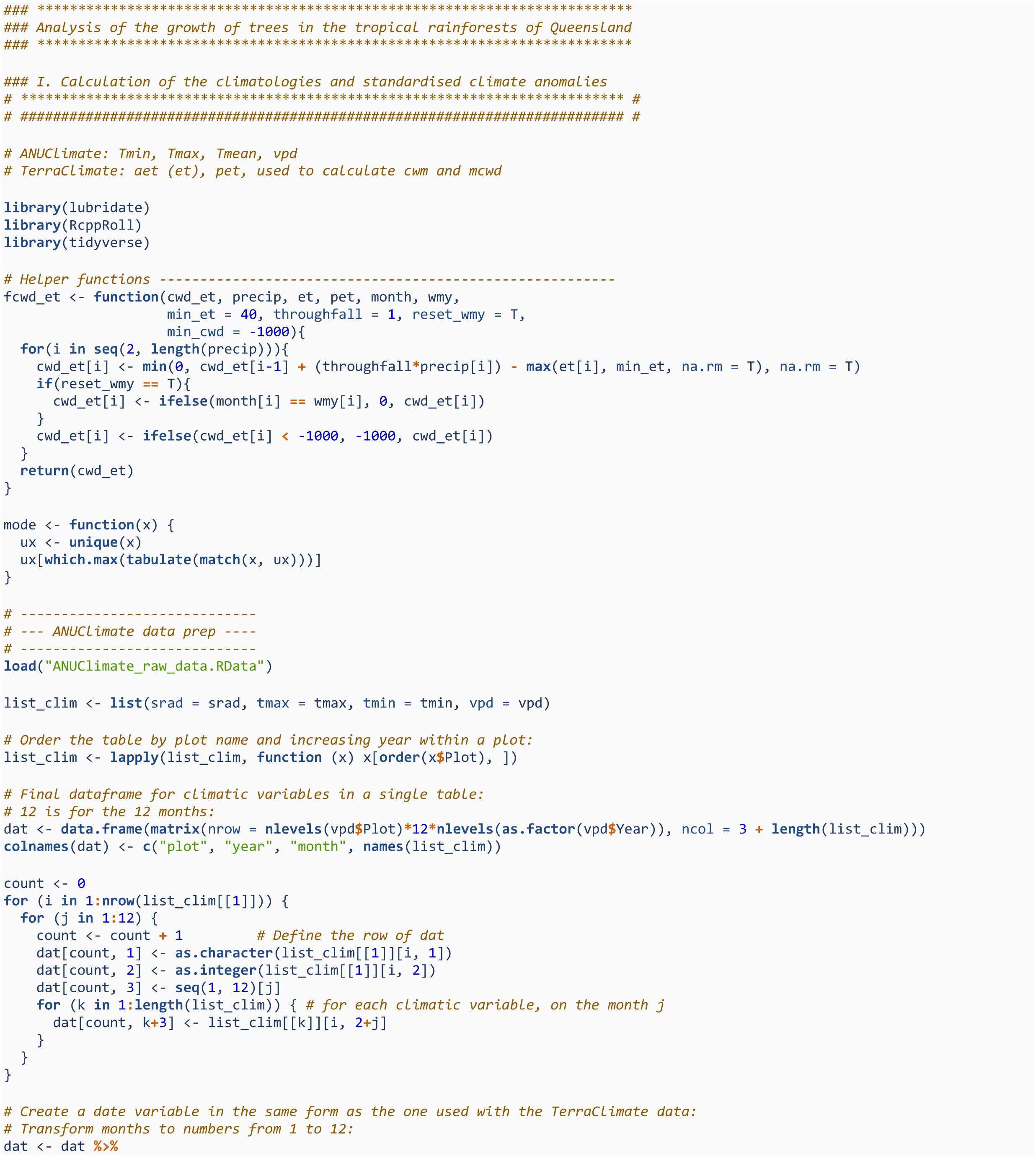
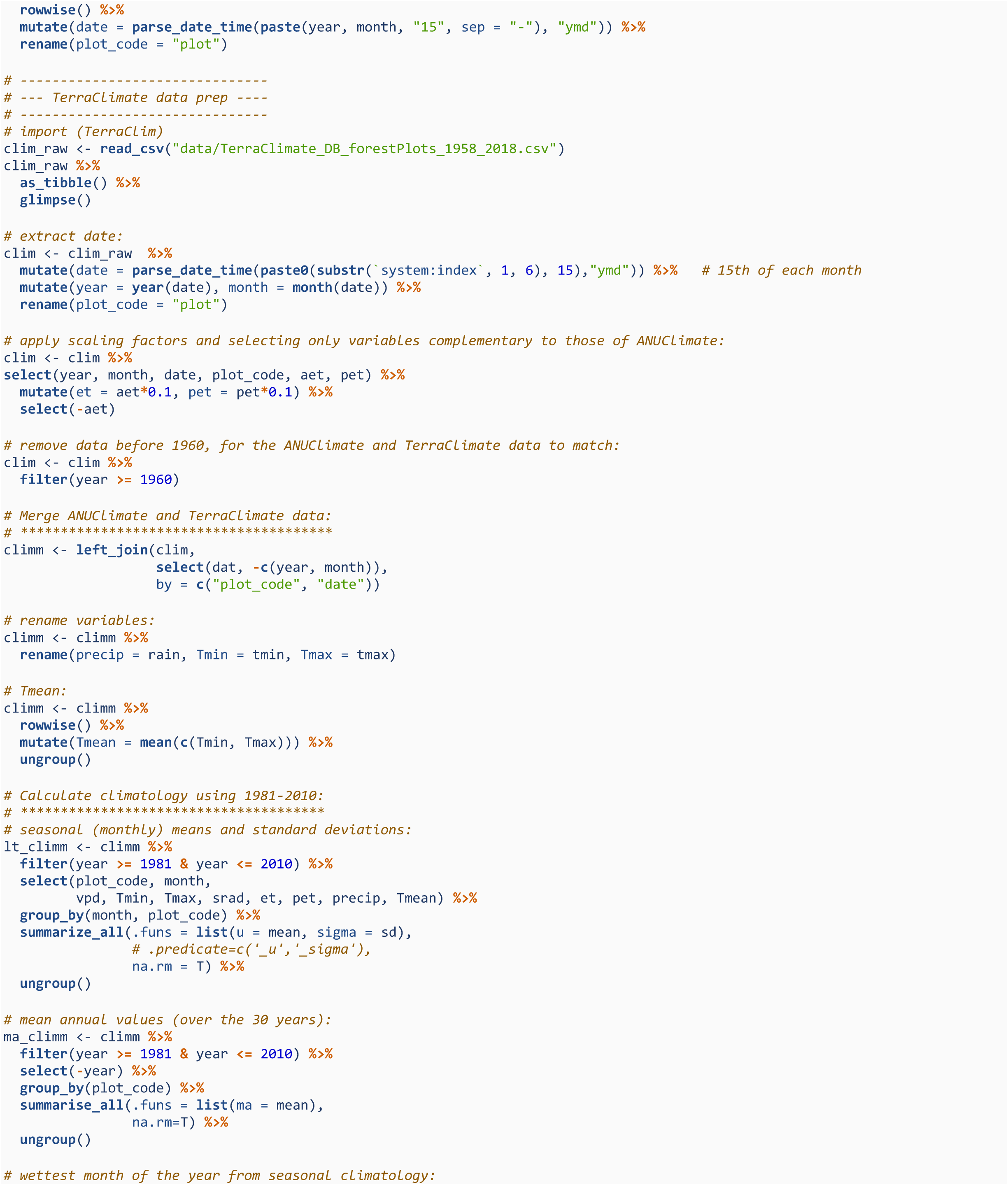
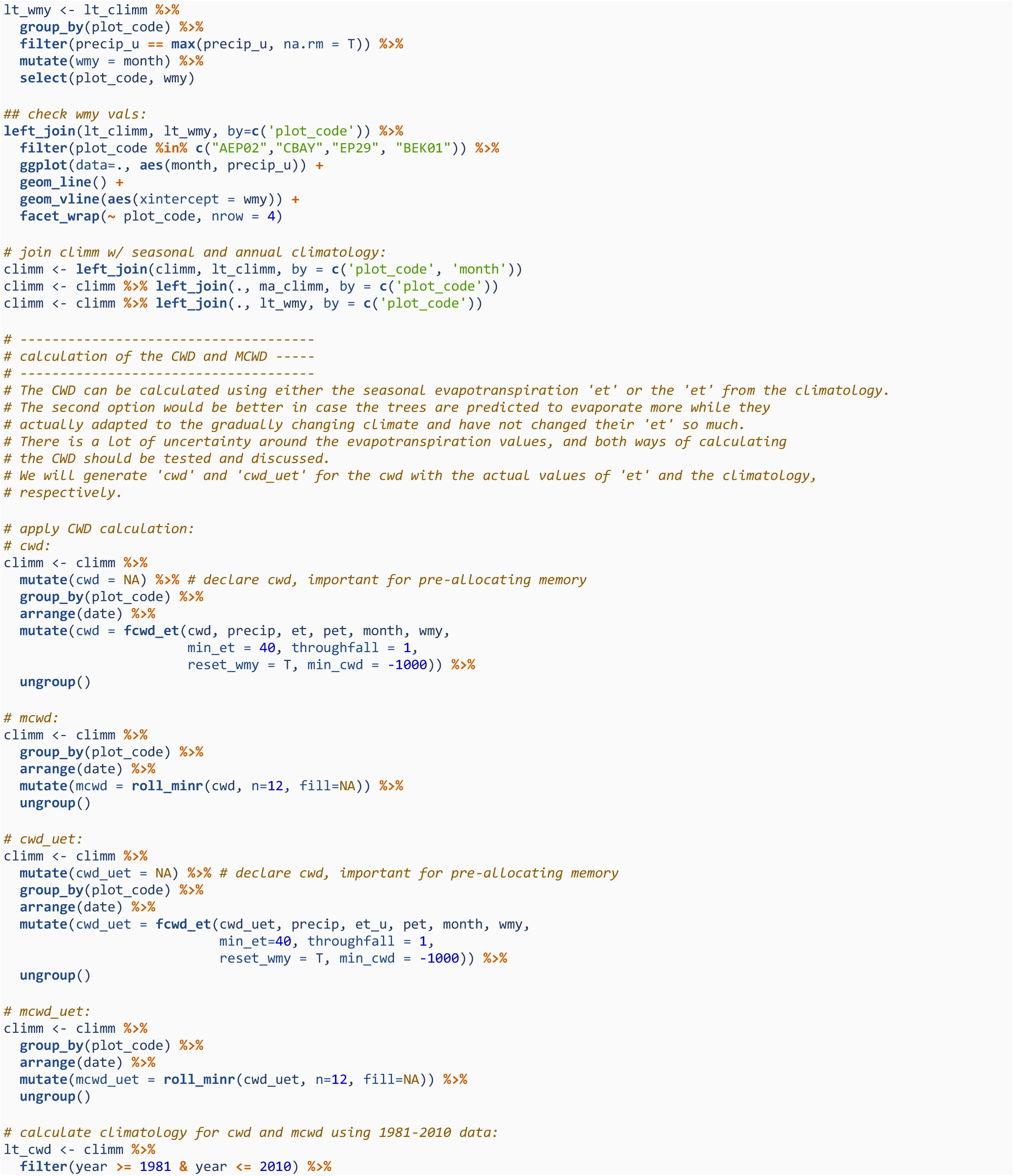
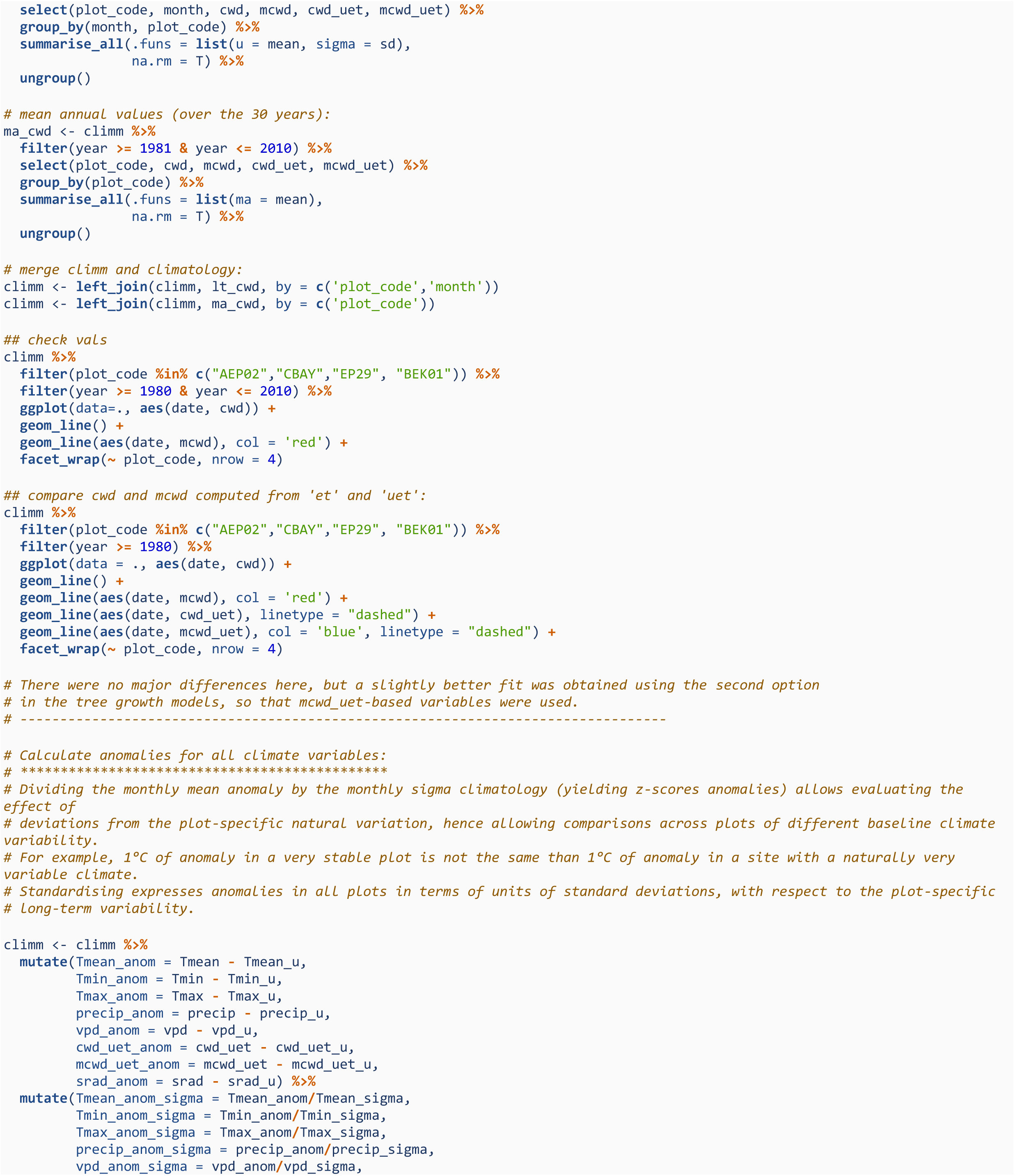
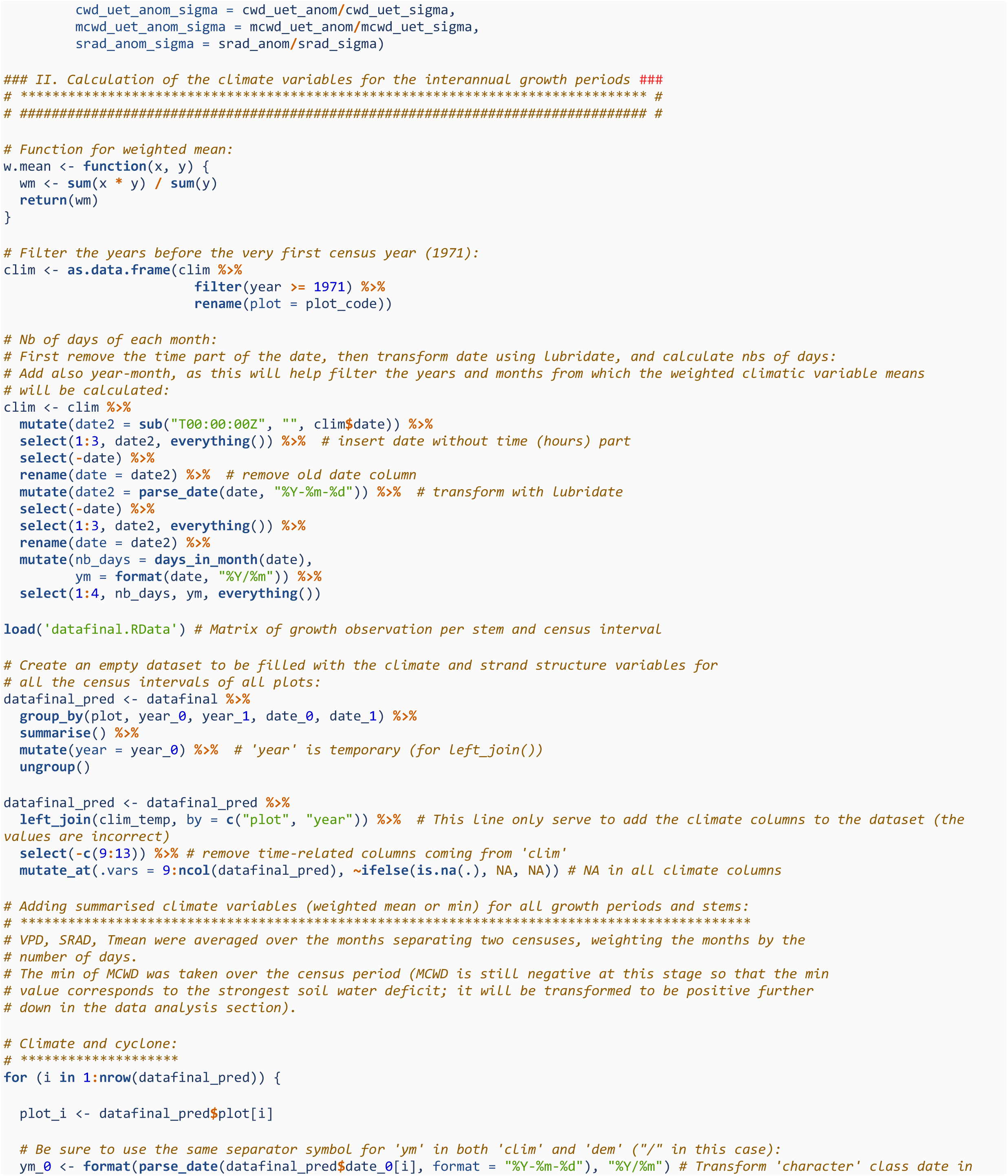
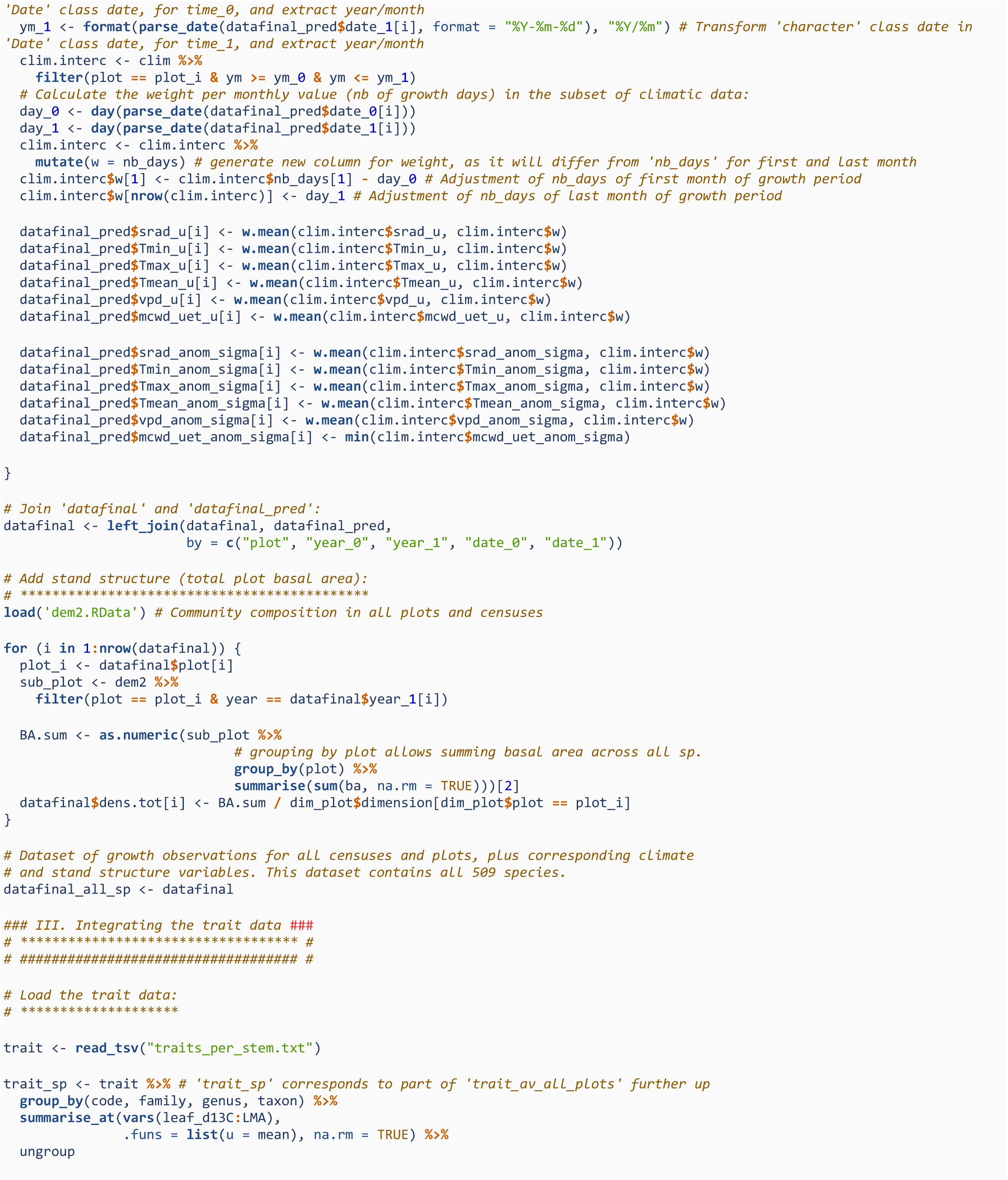
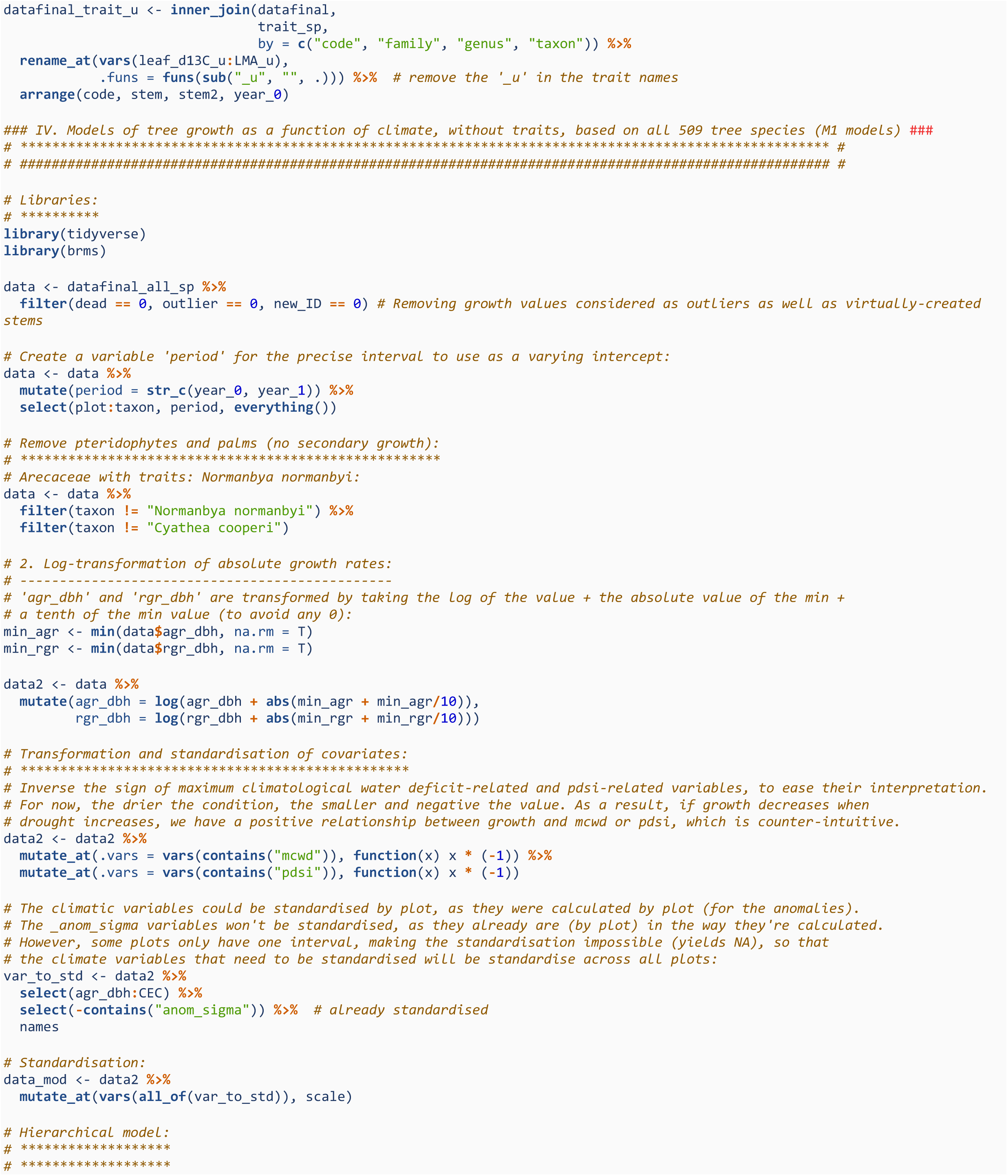
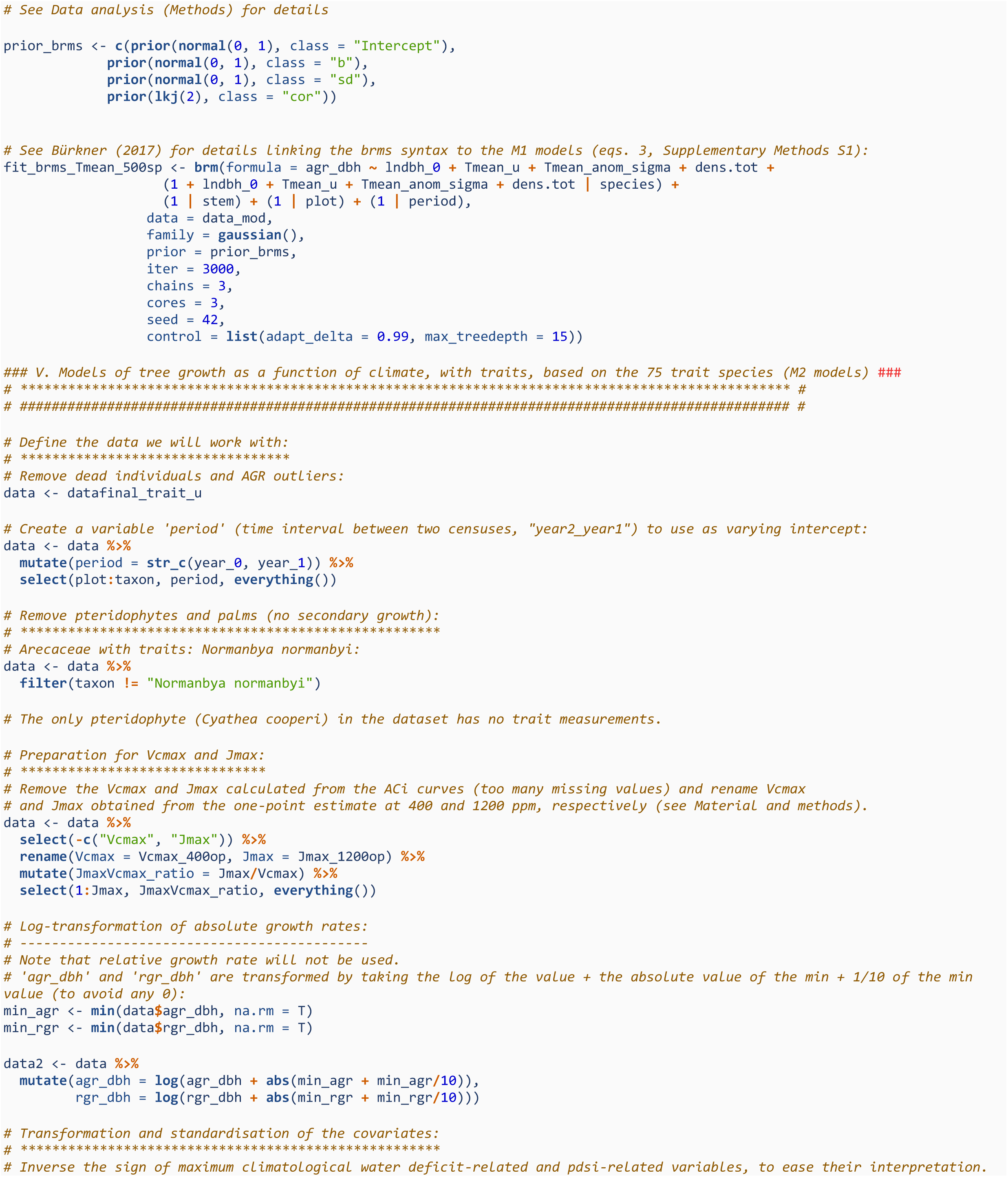
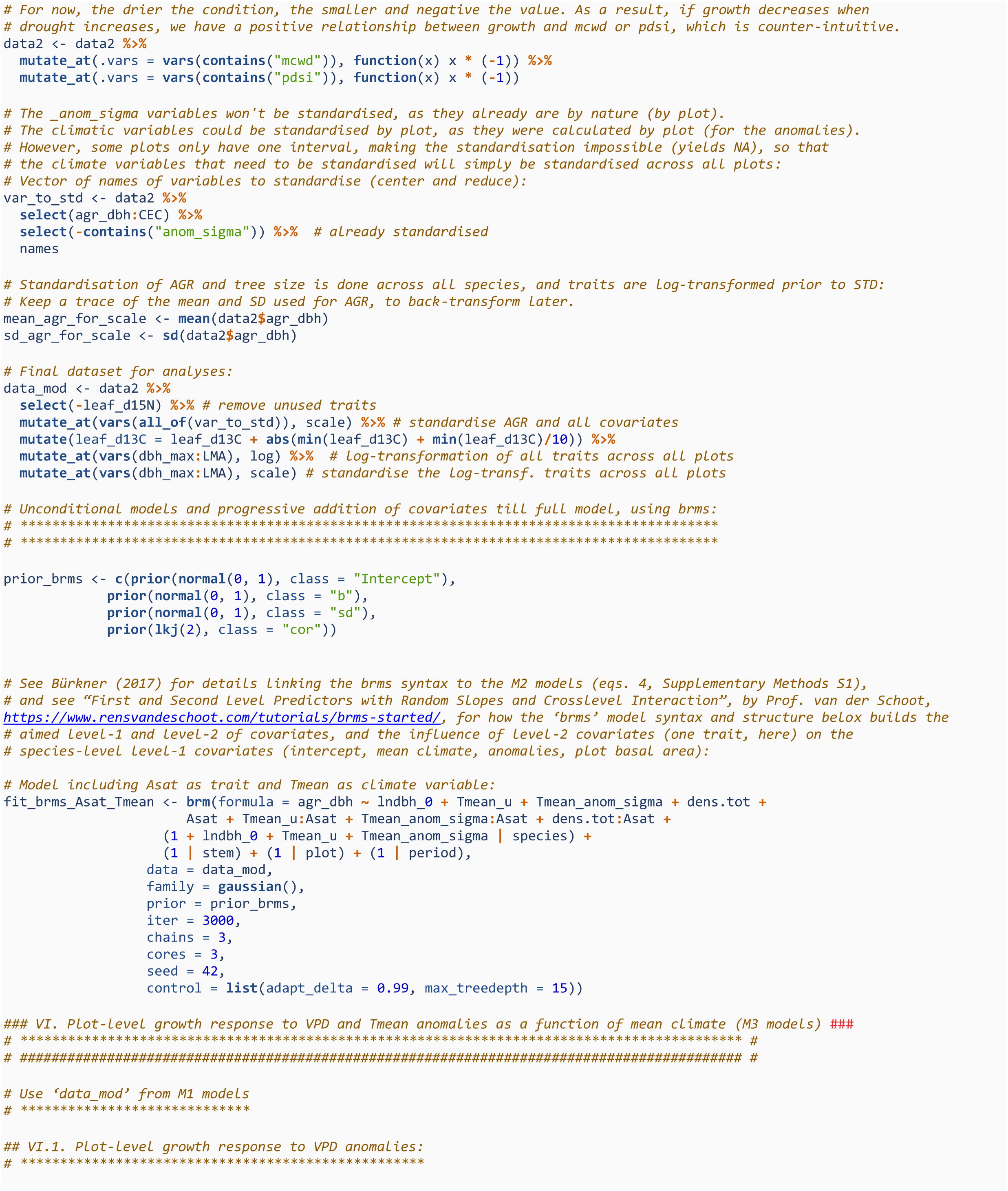
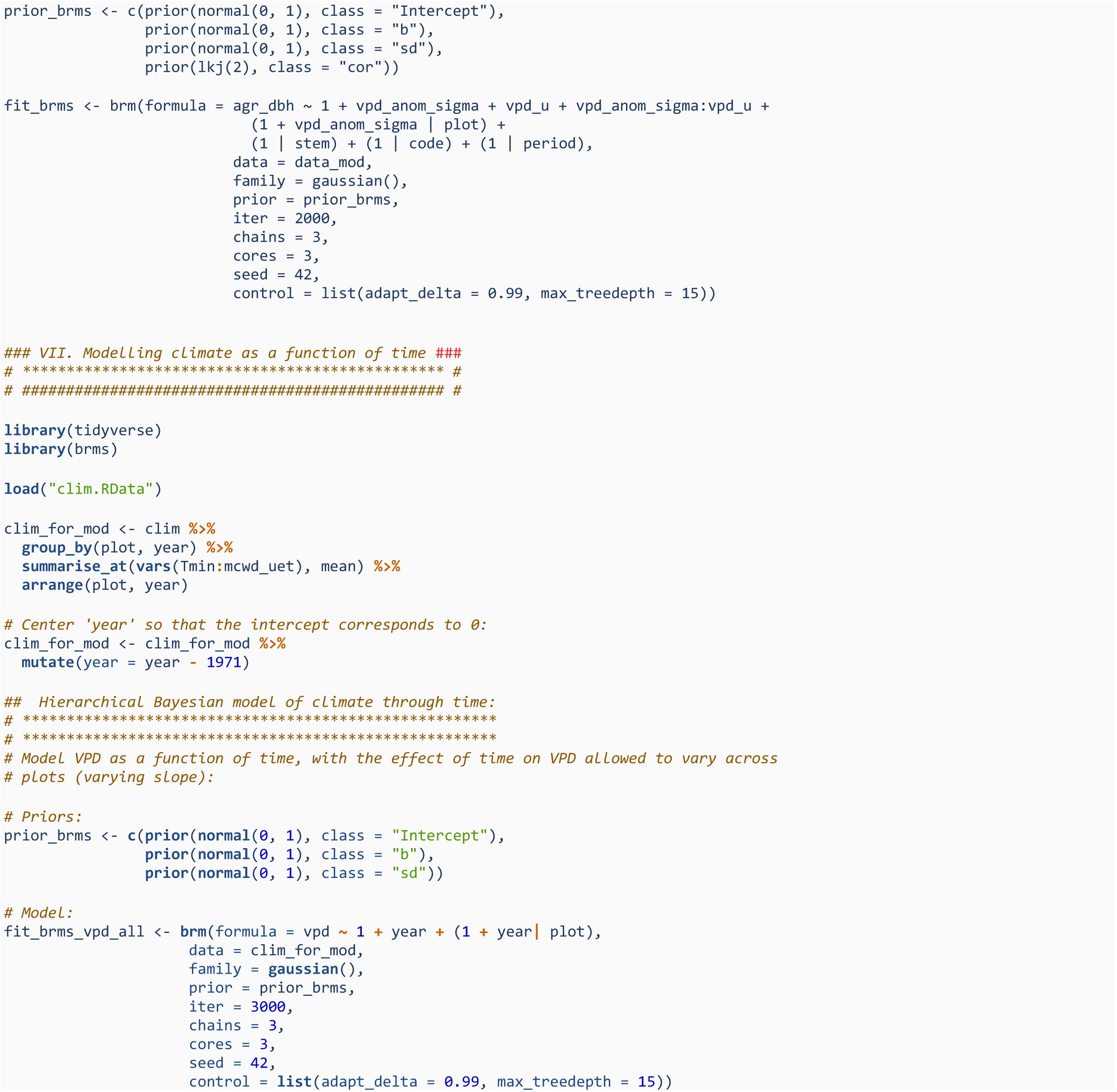

## References

1. Abatzoglou, J.T., Dobrowski, S.Z., Parks, S.A. & Hegewisch, K.C. (2018). TerraClimate, a high-resolution global dataset of monthly climate and climatic water balance from 1958-2015. Sci. Data, 5, 1–12.

2. Aguirre-Gutiérrez, J., Malhi, Y., Lewis, S.L., Fauset, S., Adu-bredu, S., Affum-Baffoe, K., et al. (2020). Long-term droughts may drive drier tropical forests towards increased functional, taxonomic and phylogenetic homogeneity. Nat. Commun., 11, 1–10.

3. Aguirre-Gutiérrez, J., Oliveras, I., Rifai, S., Fauset, S., Adu-Bredu, S., Affum-Baffoe, K., et al. (2019). Drier tropical forests are susceptible to functional changes in response to a long-term drought. Ecol. Lett., 22, 855–865.

4. Aragão, L.E.O.C., Malhi, Y., Roman-Cuesta, R.M., Saatchi, S., Anderson, L.O. & Shimabukuro, Y.E. (2007). Spatial patterns and fire response of recent Amazonian droughts. Geophys. Res. Lett., 34, L07701.

5. Atkin, O.K., Bloomfield, K.J., Reich, P.B., Tjoelker, M.G., Asner, G.P., Bonal, D., et al. (2015). Global variability in leaf respiration in relation to climate, plant functional types and leaf traits. New Phytol., 206, 614–636.

6. Bloomfield, K.J., Prentice, I.C., Cernusak, L.A., Eamus, D., Medlyn, B.E., Wright, I.J., et al. (2018). The validity of optimal leaf traits modelled on environmental conditions. New Phytol., 221, 1409–1423.

7. Bradford, M.G., Murphy, H.T., Ford, A.J., Hogan, D.L. & Metcalfe, D.J. (2014). Long-term stem inventory data from tropical rain forest plots in Australia. Ecology, 95, 2362–000.

8. Brodribb, T.J., Powers, J. & Choat, B. (2020). Hanging by a thread? Forests and drought. Science (80-.)., 368, 261–266.

9. Bürkner, P.-C. (2017). brms : An R package for Bayesian multilevel models using Stan. J. Stat. Softw., 80.

10. Carpenter, B., Gelman, A., Hoffman, M.D., Lee, D., Goodrich, B., Betancourt, M., et al. (2017). Stan: A probabilistic programming language. J. Stat. Softw., 76.

11. Cernusak, L.A., Ubierna, N., Winter, K., Holtum, J.A.M., Marshall, J.D. & Farquhar, G.D. (2013). Environmental and physiological determinants of carbon isotope discrimination in terrestrial plants. New Phytol., 200, 950–965.

12. Chave, J., Coomes, D., Jansen, S., Lewis, S.L., Swenson, N.G. & Zanne, A.E. (2009). Towards a worldwide wood economics spectrum. Ecol. Lett., 12, 351–366.

13. Choat, B., Jansen, S., Brodribb, T.J., Cochard, H., Delzon, S., Bhaskar, R., et al. (2012). Global convergence in the vulnerability of forests to drought. Nature, 491, 752–755.

14. Clark, J.S., Bell, D.M., Kwit, M.C. & Zhu, K. (2014). Competition-interaction landscapes for the joint response of forests to climate change. Glob. Chang. Biol., 20, 1979–1991.

15. Condit, R., Aguilar, S., Hernandez, A., Perez, R., Lao, S., Angehr, G., et al. (2004). Tropical forest dynamics across a rainfall gradient and the impact of an El Niño dry season. J. Trop. Ecol., 20, 51–72.

16. Condit, R., Pérez, R., Lao, S., Aguilar, S. & Hubbell, S.P. (2017). Demographic trends and climate over 35 years in the Barro Colorado 50 ha plot. For. Ecosyst., 4, 1–13.

17. Dinerstein, E., Olson, D., Joshi, A., Vynne, C., Burgess, N.D., Wikramanayake, E., et al. (2017). An ecoregion-based approach to protecting half the terrestrial realm. Bioscience, 67, 534–545.

18. Doughty, C.E. & Goulden, M.L. (2009). Are tropical forests near a high temperature threshold? J. Geophys. Res. Biogeosciences, 114, 1–12.

19. Duursma, R.A. (2015). Plantecophys - An R Package for Analysing and Modelling Leaf Gas Exchange Data. PLoS One, 10, e0143346.

20. Edwards, W., Liddell, M.J., Franks, P., Nichols, C. & Laurance, S.G.W. (2018). Seasonal patterns in rainforest litterfall: Detecting endogenous and environmental influences from long-term sampling. Austral Ecol., 43, 225–235.

21. Fadrique, B., Báez, S., Duque, Á., Malizia, A., Blundo, C., Carilla, J., et al. (2018). Widespread but heterogeneous responses of Andean forests to climate change. Nature, 564, 207–212.

22. Farquhar, G.D., Caemmerer, S. & Berry, J.A. (1980). A biochemical model of photosynthetic CO_2_ assimilation in leaves of C_3_ species. Planta, 149, 78–90–90.

23. Fortunel, C., Lasky, J.R., Uriarte, M., Valencia, R., Wright, S.J., Garwood, N.C., et al. (2018). Topography and neighborhood crowding can interact to shape species growth and distribution in a diverse Amazonian forest. Ecology, 99, 2272–2283.

24. Fortunel, C., Stahl, C., Heuret, P., Nicolini, E. & Baraloto, C. (2020). Disentangling the effects of environment and ontogeny on tree functional dimensions for congeneric species in tropical forests. New Phytol., 226, 385–395.

25. Fortunel, C., Valencia, R., Wright, S.J., Garwood, N.C. & Kraft, N.J.B. (2016). Functional trait differences influence neighbourhood interactions in a hyperdiverse Amazonian forest. Ecol. Lett., 19, 1062–1070.

26. Fyllas, N.M., Bentley, L.P., Shenkin, A., Asner, G.P., Atkin, O.K., Díaz, S., et al. (2017). Solar radiation and functional traits explain the decline of forest primary productivity along a tropical elevation gradient. Ecol. Lett., 20, 730–740.

27. Gibert, A., Gray, E.F., Westoby, M., Wright, I.J. & Falster, D.S. (2016). On the link between functional traits and growth rate: meta-analysis shows effects change with plant size, as predicted. J. Ecol., 104, 1488–1503.

28. Gray, E.F., Wright, I.J., Falster, D.S., Eller, A.S.D., Lehmann, C.E.R., Bradford, M.G., et al. (2019). Leaf:wood allometry and functional traits together explain substantial growth rate variation in rainforest trees. AoB Plants, 11, 1–11.

29. Green, J.K., Seneviratne, S.I., Berg, A.M., Findell, K.L., Hagemann, S., Lawrence, D.M., et al. (2019). Large influence of soil moisture on long-term terrestrial carbon uptake. Nature, 565, 476–479.

30. Grossiord, C., Grossiord, C., Buckley, T.N., Cernusak, L.A., Novick, K.A., Poulter, B., et al. (2020). Plant responses to rising vapor pressure deficit. New Phytol., 226, 1550–1566.

31. Harris, R.M.B., Beaumont, L.J., Vance, T.R., Tozer, C.R., Remenyi, T.A., Perkins-Kirkpatrick, S.E., et al. (2018). Biological responses to the press and pulse of climate trends and extreme events. Nat. Clim. Chang., 8, 579–587.

32. He, P., Wright, I.J., Zhu, S., Onoda, Y., Liu, H., Li, R., et al. (2019). Leaf mechanical strength and photosynthetic capacity vary independently across 57 subtropical forest species with contrasting light requirements. New Phytol., 223, 607–618.

33. Hutchinson, M.F., Kesteven, J.L. & Xu, T. (2014). Making the most of the ground based meteorological network using anomaly-based interpolation. In: Proceedings Session 5 of The Australian Energy and Water Exchange Initiative OzEWEX 2014. Canberra.

34. Hutchinson, M.F., Stein, J.L., Stein, J.A., Anderson, H. & Tickle, P.K. (2008). GEODATA 9 second DEM and D8: Digital Elevation Model Version 3 and Flow Direction Grid 2008.

35. Jentsch, A., Kreyling, J. & Beierkuhnlein, C. (2007). A new generation of climate change experiments: events, not trends. Front. Ecol. Environ., 5, 365–374.

36. De Kauwe, M.G., Lin, Y., Wright, I.J., Medlyn, B.E., Crous, K.Y., Ellsworth, D.S., et al. (2016). A test of the ‘one-point method’ for estimating maximum carboxylation capacity from field-measured, light-saturated photosynthesis. New Phytol., 210, 1130–1144.

37. Kay, M. (2020). tidybayes: Tidy Data and Geoms for Bayesian Models. R package version 2.1.1. https//mjskay.github.io/tidybayes/. DOI 10.5281/zenodo.1308151.

38. Konings, A.G., Williams, A.P. & Gentine, P. (2017). Sensitivity of grassland productivity to aridity controlled by stomatal and xylem regulation. Nat. Geosci., 10, 284–288.

39. Krause, G.H. & Winter, K. (2020). The photosynthetic system in tropical plants under high irradiance and temperature stress. In: Progress in Botany, vol. 82. pp. 131–169.

40. Laughlin, D.C., Gremer, J.R., Adler, P.B., Mitchell, R.M. & Moore, M.M. (2020). The net effect of functional traits on fitness. Trends Ecol. Evol., 1–11.

41. Liu, Z., Wu, C. & Wang, S. (2017). Predicting Forest Evapotranspiration by Coupling Carbon and Water Cycling Based on a Critical Stomatal Conductance Model. IEEE J. Sel. Top. Appl. Earth Obs. Remote Sens., 10, 4469–4477.

42. Malhi, Y., Aragao, L.E.O.C., Galbraith, D., Huntingford, C., Fisher, R., Zelazowski, P., et al. (2009). Exploring the likelihood and mechanism of a climate-change-induced dieback of the Amazon rainforest. Proc. Natl. Acad. Sci., 106, 20610–20615.

43. Malhi, Y., Doughty, C.E., Goldsmith, G.R., Metcalfe, D.B., Girardin, C.A.J., Marthews, T.R., et al. (2015). The linkages between photosynthesis, productivity, growth and biomass in lowland Amazonian forests. Glob. Chang. Biol., 21, 2283–2295.

44. McElreath, R. (2020). Statistical rethinking: A Bayesian course with examples in R and Stan. CRC Press, Abingdon.

45. McGill, B.J., Enquist, B.J., Weiher, E. & Westoby, M. (2006). Rebuilding community ecology from functional traits. Trends Ecol. Evol., 21, 178–185.

46. Mendivelso, H.A., Camarero, J.J., Gutiérrez, E. & Zuidema, P.A. (2014). Time-dependent effects of climate and drought on tree growth in a Neotropical dry forest: Short-term tolerance vs. long-term sensitivity. Agric. For. Meteorol., 188, 13–23.

47. Muledi, J., Bauman, D., Jacobs, A., Meerts, P., Shutcha, M. & Drouet, T. (2020). Tree growth, recruitment, and survival in a tropical dry woodland: The importance of soil and functional identity of the neighbourhood. For. Ecol. Manage., 460, 117894.

48. Murphy, H.T., Bradford, M.G., Dalongeville, A., Ford, A.J. & Metcalfe, D.J. (2013). No evidence for long-term increases in biomass and stem density in the tropical rain forests of Australia. J. Ecol., 101, 1589–1597.

49. Needham, J., Merow, C., Chang-Yang, C.-H., Caswell, H. & McMahon, S. (2018). Inferring forest fate from demographic data: from vital rates to population dynamic models. Proc. R. Soc. B, 285, 20172050.

50. Novick, K.A., Ficklin, D.L., Stoy, P.C., Williams, C.A., Bohrer, G., Oishi, A.C., et al. (2016). The increasing importance of atmospheric demand for ecosystem water and carbon fluxes. Nat. Clim. Chang., 6, 1023–1027.

51. Osnas, J.L.D., Lichstein, J.W., Reich, P.B. & Pacala, S.W. (2013). Global leaf trait relationships: Mass, area, and the leaf economics spectrum. Science (80-.)., 340, 741–744.

52. Paine, C.E.T., Amissah, L., Auge, H., Baraloto, C., Baruffol, M., Bourland, N., et al. (2015). Globally, functional traits are weak predictors of juvenile tree growth, and we do not know why. J. Ecol., 103, 978–989.

53. Pan, Y., Birdsey, R.A., Fang, J., Houghton, R., Kauppi, P.E., Kurza, W.A., et al. (2011). A large and persistent carbon sink in the world’s forest. Science (80-.)., 333, 988–993.

54. Phillips, O.L., Aragão, L.E.O.C., Lewis, S.L., Fisher, J.B., Lloyd, J., López-González, G., et al. (2009). Drought sensitivity of the amazon rainforest. Science (80-.)., 323, 1344–1347.

55. Poorter, L., Wright, S.J., Paz, H., Ackerly, D.D., Condit, R., Ibarra-Manríquez, G., et al. (2008). Are functional traits good predictors of demographic rates? Evidence from five neotropical forests. Ecology, 89, 1908–1920.

56. Powers, J.S., Vargas G., G., Brodribb, T.J., Schwartz, N.B., Pérez-Aviles, D., Smith-Martin, C.M., et al. (2020). A catastrophic tropical drought kills hydraulically vulnerable tree species. Glob. Chang. Biol., 26, 3122–3133.

57. Quebbeman, J.A. & Ramirez, J.A. (2016). Optimal allocation of leaf-level nitrogen: Implications for covariation of Vcmax and Jmax and photosynthetic downregulation. J. Geophys. Res. Biogeosciences, 121, 2464–2475.

58. Reich, P.B. (2014). The world-wide ‘fast–slow’ plant economics spectrum: a traits manifesto. J. Ecol., 102, 275–301.

59. Rifai, S.W., Girardin, C.A.J., Berenguer, E., Del Aguila-Pasquel, J., Dahlsjö, C.A.L., Doughty, C.E., et al. (2018). ENSO Drives interannual variation of forest woody growth across the tropics. Philos. Trans. R. Soc. B Biol. Sci., 373.

60. Rifai, S.W., Li, S. & Malhi, Y. (2019). Coupling of El Niño events and long-term warming leads to pervasive climate extremes in the terrestrial tropics. Environ. Res. Lett., 14.

61. Roeber, V.M., Bajaj, I., Rohde, M., Schmülling, T. & Cortleven, A. (2020). Light acts as a stressor and influences abiotic and biotic stress responses in plants. Plant. Cell Environ., pce.13948.

62. Rosas, T., Martínez-Vilalta, J., Mencuccini, M., Cochard, H., Barba, J. & Saura-Mas, S. (2019). Adjustments and coordination of hydraulic, leaf and stem traits along a water availability gradient. New Phytol., 223, 632–646.

63. Rowland, L., Oliveira, R.S., Bittencourt, P.R.L., Giles, A.L., Coughlin, I., Costa, P. de B., et al. (2021). Plant traits controlling growth change in response to a drier climate. New Phytol., 229, 1363–1374.

64. Rüger, N., Wirth, C., Wright, S.J. & Condit, R. (2012). Functional traits explain light and size response of growth rates in tropical tree species. Ecology, 93, 2626–2636.

65. Sánchez-Salguero, R., Linares, J.C., Camarero, J.J., Madrigal-González, J., Hevia, A., Sánchez-Miranda, Á., et al. (2015). Disentangling the effects of competition and climate on individual tree growth: A retrospective and dynamic approach in Scots pine. For. Ecol. Manage., 358, 12–25.

66. Sanginés de Cárcer, P., Vitasse, Y., Peñuelas, J., Jassey, V.E.J., Buttler, A. & Signarbieux, C. (2018). Vapor–pressure deficit and extreme climatic variables limit tree growth. Glob. Chang. Biol., 24, 1108–1122.

67. Schielzeth, H. (2010). Simple means to improve the interpretability of regression coefficients. Methods Ecol. Evol., 1, 103–113.

68. Smith, N.G., Keenan, T.F., Colin Prentice, I., Wang, H., Wright, I.J., Niinemets, Ü., et al. (2019). Global photosynthetic capacity is optimized to the environment. Ecol. Lett., 22, 506–517.

69. Sullivan, M.J.P., Lewis, S.L., Affum-Baffoe, K., Castilho, C., Costa, F., Sanchez, A.C., et al. (2020). Long-term thermal sensitivity of Earth’s tropical forests. Science (80-.)., 368, 869–874.

70. Team, R.C. (2020). R: A Language and Environment for Statistical Computing. R Foundation for Statistical Computing, Vienna, Austria.

71. Tjoelker, M.G., Oleksyn, J. & Reich, P.B. (2001). Modelling respiration of vegetation: Evidence for a general temperature-dependent Q10. Glob. Chang. Biol., 7, 223–230.

72. Uriarte, M., Lasky, J.R., Boukili, V.K. & Chazdon, R.L. (2016). A trait-mediated, neighbourhood approach to quantify climate impacts on successional dynamics of tropical rainforests. Funct. Ecol., 30, 157–167.

73. Vile, D., Garnier, É., Shipley, B., Laurent, G., Navas, M.L., Roumet, C., et al. (2005). Specific leaf area and dry matter content estimate thickness in laminar leaves. Ann. Bot., 96, 1129–1136.

74. Violle, C., Navas, M.-L., Vile, D., Kazakou, E., Fortunel, C., Hummel, I., et al. (2007). Let the concept of trait be functional! Oikos, 116, 882–892.

75. Wagner, F., Rossi, V., Baraloto, C., Bonal, D., Stahl, C. & Hérault, B. (2014). Are commonly measured functional traits involved in tropical tree responses to climate? *Int*. J. Ecol., 2014, 389409.

76. Walker, A.P., Beckerman, A.P., Gu, L., Kattge, J., Cernusak, L.A., Domingues, T.F., et al. (2014). The relationship of leaf photosynthetic traits – Vcmax and Jmax – to leaf nitrogen, leaf phosphorus, and specific leaf area: a meta-analysis and modeling study. Ecol. Evol., 4, 3218–3235.

77. Westoby, M. (1998). A leaf-height-seed (LHS) plant ecology strategy scheme. Plant Soil, 199, 213–227.

78. Westoby, M., Falster, D.S., Moles, A.T., Vesk, P.A. & Wright, I.J. (2002). Plant ecological strategies: Some leading dimensions of variation between species. Annu. Rev. Ecol. Syst., 33, 125–159.

79. Wickham, H., Averick, M., Bryan, J., Chang, W., McGowan, L., François, R., et al. (2019). Welcome to the tidyverse. J. Open Source Softw., 4, 1686.

80. Wills, J., Herbohn, J., Hu, J., Sohel, S., Baynes, J. & Firn, J. (2018). Tree leaf trade-offs are stronger for sub-canopy trees: leaf traits reveal little about growth rates in canopy trees. Ecol. Appl., 28, 1116–1125.

81. Wright, I.J., Dong, N., Maire, V., Prentice, I.C., Westoby, M., Díaz, S., et al. (2017). Global climatic drivers of leaf size. Science (80-.)., 357, 917–921.

82. Wright, I.J., Reich, P.B., Westoby, M., Ackerly, D.D., Baruch, Z., Bongers, F., et al. (2004). The worldwide leaf economics spectrum. Nature, 428, 821–827.

83. Yuan, W., Zheng, Y., Piao, S., Ciais, P., Lombardozzi, D., Wang, Y., et al. (2019). Increased atmospheric vapor pressure deficit reduces global vegetation growth. Sci. Adv., 5, 1–13.

